# Crossover model of Lep-Rec reveals higher heritability of recombination

**DOI:** 10.1101/2024.12.23.630188

**Authors:** Pasi Rastas, Teemu Kivioja

## Abstract

Meiotic recombination, a process that reshuffles DNA between parental chromosomes, is almost universal in multicellular organisms. Recombination accelerates the response to selection by breaking the linkage and creating new allelic combinations that can affect the fitness of the progeny. This motivates us to characterise recombination rate variation and to take it into account in evolutionary models and studies.

Linkage mapping utilises recombination to obtain recombination distances for genetic markers. With (whole) genome sequencing data, very dense linkage maps can be produced, locating crossovers precisely in the genome. This enables direct and accurate calculation of recombination distances, correction of possible errors in the genome and maps, and studying the relation between recombination and physical base-pair distances. This is now a relevant problem, as high-quality genomes are emerging for many species, and available dense linkage map data would supplement these genomes.

Here we present a novel software Lep-Rec to compute the local recombination rate, i.e. the percentage of crossovers per individual per megabase (cM/Mb) along the genome. Moreover, it can also estimate the underlying, only partly observed, tetrad crossover distribution for each chromosome, while modelling crossover and chromatid interference. Together with Lep-MAP3 and Lep-Anchor, Lep-Rec forms a complete toolbox for studying recombination and crossovers: Lep-MAP3 can robustly construct linkage maps for large number of markers and individuals, while Lep-Anchor can anchor, validate and correct genome assemblies using linkage maps, and together these software provide consistent and complete physical and linkage maps for further analysis with Lep-Rec. Lep-Rec is available from http://sourceforge.net/projects/lep-anchor.

Finally, we demonstrate the performance of Lep-Rec using real and simulated data: It outperforms and simplifies currently available tools and its estimated crossover distribution can improve association analysis and heritability estimates of recombination.

## 1 Introduction

Two genetic markers within a chromosome are one centimorgan (cM) apart if the expected number of observed crossovers between them is 1% within one gamete (chromatid) and generation (Sturtevant (1913)). If the crossover locations are known for some population sample, the cM distance between any two markers can be directly computed from the number of crossovers between them. Moreover, the local recombination rate (cM/Mb) can be defined as the centimorgan distance of a genomic region divided by the region’s length in megabases (Mb). We assume that a genome assembly defines the genomic regions and physical locations of the markers.

A Marey map (Chakravarti, 1991) is a line chart of the physical position (in Mb) to the linkage position (in cM). It is a common way to visualise the relation between the physical and genetic (linkage map) distances. The local recombination rate (cM/Mb), can be obtained as the slope (derivative) of the Marey map, assuming the physical position is the *x*-axis. Conversely, Marey map can be obtained as the cumulative function (integral) of the recombination rate. Recombination rates and Marey maps are complementary; A Marey map emphasises the total map length while the physical distribution of crossovers is more apparent from a recombination rate (see Figure 5). With sex-specific recombination rates and Marey maps, we can detect sex differences in recombination rates (heterochiasmy) or landscapes (Sardell and Kirkpatrick, 2020).

With data on a suitable cross, e.g. outbred F1, F2 or backcross, linkage mapping locates paternal and maternal crossovers (between two markers) for each offspring (Rastas, 2017), in addition to the linkage (cM) positions of the markers. These sex-specific crossover locations enable inferences of the four parental meiotic products, of which an individual inherits a gamete (also called single spore (Zhao et al., 1995)) from both parents. This analysis of tetrad crossovers is explained in more detail in Figure 4. With the tetrad analysis, we can study phenomena like 1) crossover interference, where each crossover reduces the probability of a neighbouring crossover (see Otto and Payseur (2019)), 2) chromatid interference (Zhao et al., 1995; Sarens et al., 2021), possible biases in the participation of non-sister chromatid pairs in crossovers (see Figure 4) and 3) obligatory crossover (or crossover assurance) (Wang et al., 2015).

The linkage positions might be inconsistent with the physical positions, due to errors and uncertainty (Rastas, 2020). In order to study recombination, these inconsistencies must be resolved first. In this article, we study how to make the linkage and physical maps consistent, and then, how to estimate recombination rates, Marey maps and tetrad crossovers from consistent maps.

This is now a relevant problem as the number of species with high-quality genome assemblies is increasing due to better data and data analysis (e.g. Cheng et al. (2021)), and for several species there are new and old linkage mapping data to supplement these genomes, e.g. Stapley et al. (2017) found linkage mapping results for 353 species. In addition, the linkage mapping software and the corresponding data analysis have recently been improved to extract more information from the available data (Rastas, 2017). We assume here that linkage maps are very dense, e.g. constructed from (whole) genome sequencing data. Then most, if not all, crossovers can be located between two close-by markers in the genome, and the map positions can be calculated directly from the number of crossovers without a mapping function (see Peñalba and Wolf (2020) about mapping functions).

### 1.1 Relation to previous work

A typical way to estimate the local recombination is to fit some smooth polynomial (spline) function to a scatter plot of the physical and linkage positions of the markers, and then differentiate this function (Siberchicot et al., 2017; Rezvoy et al., 2007). However, fitted polynomial function could be decreasing, making the local recombination rate negative (rate should be ≥ 0). An example of negative recombination rates from a spline fit is given in Figure 5. We also noticed this problem using MareyMap online software (Siberchicot et al., 2017) (data not shown). Moreover, a simple fit cannot correct or ignore possible erroneous linkage positions.

Our non-parametric method Lep-Rec solves these problems by fitting a piece-wise constant and non-decreasing step function to the (noisy) data and extracting (possibly corrected) crossovers between the marker positions. Then variable kernel density estimation (see Terrell and Scott (1992)) is used to obtain a smooth and non-negative recombination rate from the crossover locations. The obtained recombination rate can be integrated back into a continuous, non-decreasing and smoothed Marey map. This fitting is applicable to data with only marker positions (see Figure 2 (b)), e.g. enabling re-analysis of Marey maps extracted from the literature. Lep-Rec does its computation in an automatic fashion and, in most cases, without any manual steps, whereas using available software, e.g. MareyMap online (Siberchicot et al., 2017), requires tedious manual work for every chromosome and map.

The local recombination rate can also be estimated from genetic population data or from gamete sequencing (Peñalba and Wolf, 2020; Chen et al., 2022). These methods only produce average or single-sex maps without full information on the gametes and require high-quality genome assembly. However, the linkage map data, used by Lep-Rec, enables error correction of the genome and extracts sex-specific gametes and recombination rates.

The tetrad crossover distribution model was originally proposed by Weinstein (1936) and its maximum likelihood estimation by Yu and Feingold (2001) and by Ott (1996), but also used in the count-location model (see McPeek and Speed (1995)) originally by Karlin and Liberman (1979). A simple model for the crossover locations (in cM) without interference is the Poisson point process with a rate parameter *λ* (McPeek and Speed, 1995). The number of crossovers in this model is Poisson distributed and the distance of adjacent crossovers (as well as the distance of the first and the last crossover to the chromosome start and end, respectively) is exponentially distributed with parameter *λ*. However, conditional on the number of crossovers in the relative chromosome position [0, 1], the distances are uniformly distributed.

The gamma model has been proposed as a generalisation of the Poisson process with crossover interference (McPeek and Speed, 1995). There, the distance of adjacent crossovers *d* ∼is gamma distributed as *d* gamma(*ν, λ · ν*). When *ν* = 1 there is no interference and this corresponds to the Poisson rocess with rate *λ*. Typically, only positive crossover interference is considered, i.e. *ν* ≥ 1. Further generalisation is the gamma sprinkling model (Copenhaver et al., 2002; Housworth and Stahl, 2003), where the distances are distributed as a mixture of gamma and exponential distributions, and the parameter *p* is the mixture fraction of the exponential distribution. The parameter *ν* has been used to quantify crossover interference, e.g. listed by (Otto and Payseur, 2019). There are also many other mechanical and statistical crossover models, see Otto and Payseur (2019) for a more complete listing of them. We have tried two existing software using the gamma (sprinkling) model, CODA (Gauthier et al., 2011) and the software of Campbell et al. (2015). These software produced similar parameter values for *λ* and *ν* as our experiments with the same models and data.

Here we extend the method of Yu and Feingold (2001) to take into account the physical locations of crossovers, crossover and chromatid interference (studied earlier by Zhao et al. (1995)) and errors in the data. The main difference is that we model the physical positions of crossovers, whereas most previous works (e.g McPeek and Speed (1995)) are modelling the genetic (cM) positions. Our software is also fast, easy to use, and it implements these new and existing gamma (sprinkling) models and their extensions with obligatory crossover, chromatid interference and data error models.

## 2 Methods

The workflow of Lep-Rec is depicted in Figure 1. The main module is EstimateRecombination for estimating recombination rates and conducting tetrad analysis. Its input consists of one or several linkage map(s) with consistent physical and linkage positions. The input data can be filtered on, e.g. the number of supporting markers or the number of crossovers per individual.

**Figure 1:**
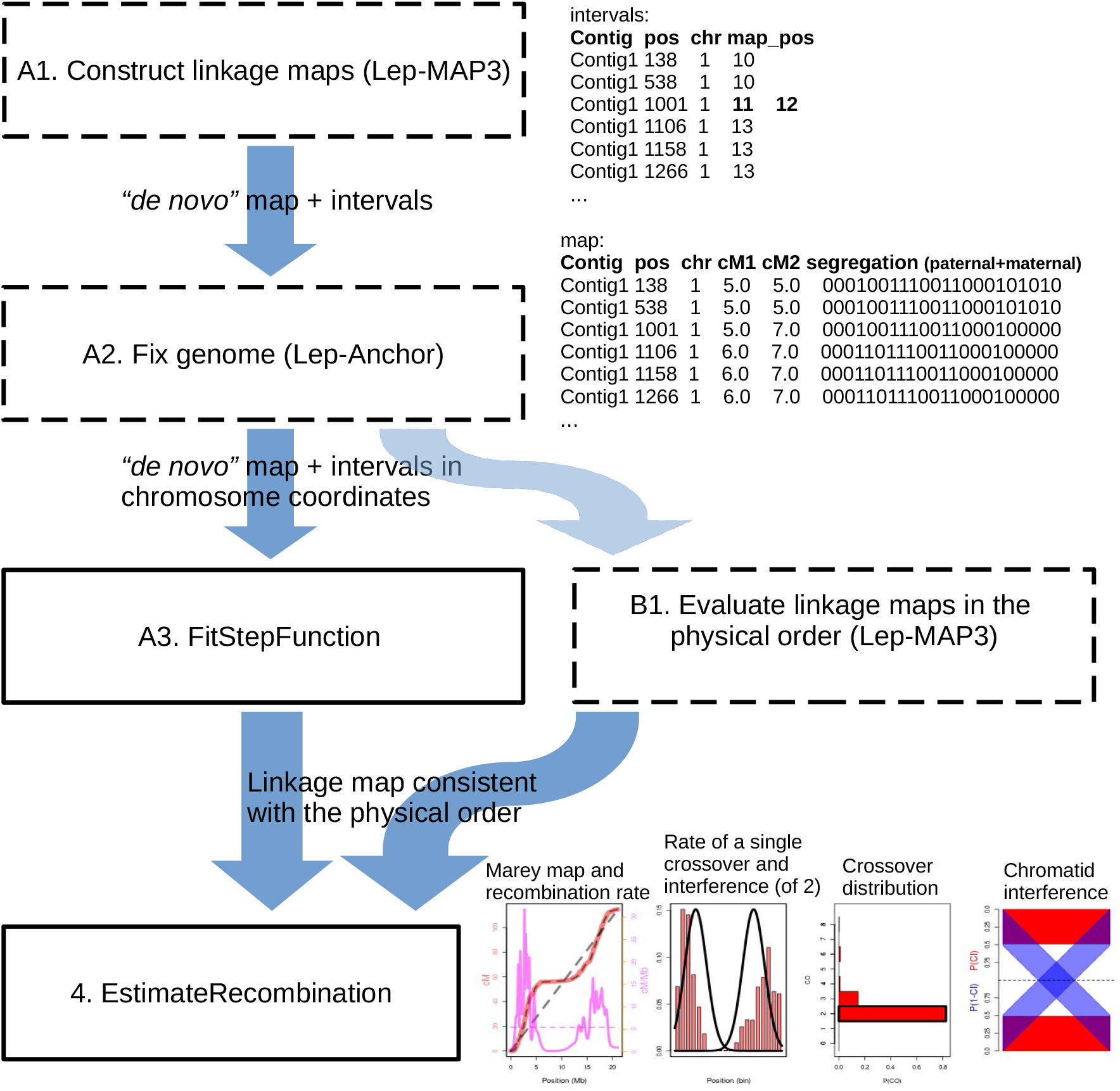
The workflow of Lep-Rec to obtain recombination estimates from linkage map(s) in the genome coordinates.

If the maps are constructed *de novo* (independently of the genome), the module FitStepFunction can be used to correct maps to be consistent with the physical marker order. Another option is to order the genetic markers into the (correct) physical order and evaluate the linkage map assuming this marker order. Such map evaluation is possible with Lep-MAP3 (Rastas, 2017), via parameter evaluateOrder in OrderMarkers2.

### 2.1 FitStepFunction

FitStepFunction module fits a piecewise constant and non-decreasing step function to the scatter plot of the physical and linkage positions (Figure 2 (a)). To cope with obscure map positions due to noise and uncertainty, FitStepFunction uses the same algorithm as Lep-Anchor (Rastas, 2020) uses to calculate the anchoring score to place contigs within a chromosome. The input is a linkage map with markers in the physical coordinates and optionally marker position intervals (from Lep-MAP3) to account for the uncertainty in the map data. The fitted step function anchors each marker to a map position, and this position is output together with the corresponding segregation pattern(s). The segregation patterns are later used to detect the crossover locations (Figure 2 (c)). When necessary, this module also reverses the linkage map to the same orientation as the physical map, and trims the map to start at 0 cM.

**Figure 2:**
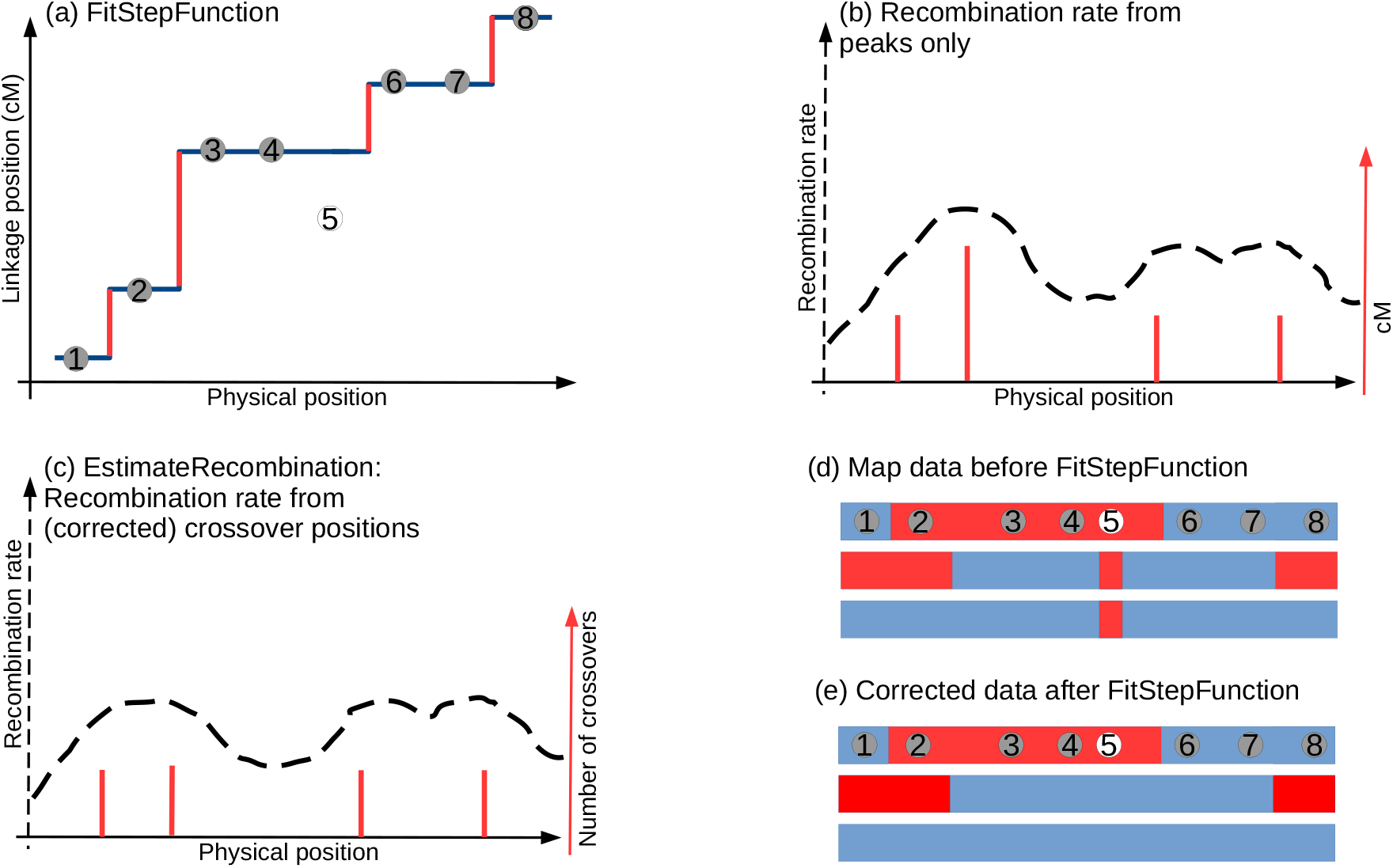
Module FitStepFunction and recombination rate. (a) The FitStepFunction fits a piecewise constant non-decreasing step function to the linkage positions of the markers in physical order. The fit is maximising the number of consistent markers (7 out of 8). The corresponding steps indicating crossovers are shown in red. (b) These steps can be used to estimate the local recombination rate (dashed line) as density estimation. This always yields a non-negative rate. (c) After the FitStepFunction, crossovers supported only by the inconsistent marker 5 have been removed and the EstimateRecombination estimates recombination rate from the remaining crossover positions. (d) The map data (haplotypes) before FitStepFunction and after (e), the color changes indicate crossovers.

### 2.2 EstimateRecombination: local recombination rate

First, module EstimateRecombination calculates smoothed recombination rate. It handles sparse data by smoothing and uncertain crossover location by sampling 5 (parameter numSamples) locations from each position interval between corresponding markers. The intervals are extracted individually from the segregation patterns of the markers. By default, variable kernel density estimation (see Terrell and Scott (1992)) is used to calculate the local recombination rate from the sampled locations. There, Gaussian kernels are added to each crossover location *l* with the standard deviation as the distance between *l* and the *S*:th nearest crossover to it. The parameter *S* controlling the local smoothness, can be given by the user or obtained automatically by a (five-fold) cross-validation procedure. Instead of these variable kernels, a constant deviation (e.g. 2 Mb) can be provided as well. Finally, the obtained recombination rate can be integrated back into a continuous and smooth Marey map. As the density is naturally non-negative, the obtained Marey map will be non-decreasing.

### 2.3 EstimateRecombination: tetrad crossover analysis

After estimating the local recombination rate, EstimateRecombination proceeds with the tetrad analysis, i.e. it classifies the individuals based on the number of underlying tetrad crossovers, of which only half, on average, are detected from the data. It extends the maximum likelihood method of Yu and Feingold (2001) with the ideas of Zhao et al. (1995) into a complete and computationally feasible solution. These analyses require data where maternal and paternal crossovers can be separated, e.g. outbred F1, F2 or backcross.

Consider that the data consist of a sample of *M* gametes from *M* independent paternal or maternal meioses. The observed crossovers in each gamete can be summarised by a vector ***n*** = (*n*_0_, *n*_1_, …, *n*_*N*_), where *n*_*i*_ is the number of gametes with *i* observed crossovers and *N* is the maximum number of crossovers in the sample. In the Yu and Feingold (2001), the likelihood of data ***n*** is

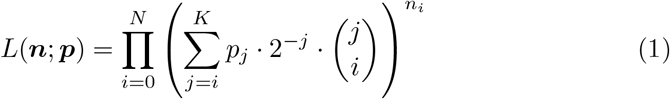

where *K* is the maximum number of tetrad crossovers (*N* ≤ *K* ≤ 2*N*), ***p*** = (*p*_0_, *p*_1_, …, *p*_*K*_) is a vector where *p*_*j*_ is the probability (or rate) of having exactly *j* crossovers in a tetrad. The free parameters *p*_*j*_ are obtained by maximising the likelihood of (1) using an EM algorithm. Note that this method ignores crossover positions and chromatid interference.

In Lep-Rec, we summarise the recombination data by an *M* -tuple ***r*** = (*r*_1_, *r*_2_, …, *r*_*M*_), where each *r*_*i*_ is the set of observed crossover positions for the gamete *i*. The generalised data likelihood is

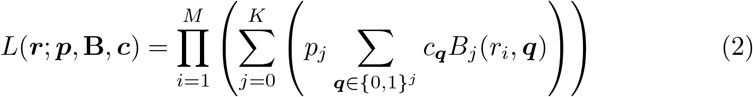

where ***p*** and *K* are as in (1), ***q*** is the crossover pattern indicating which of the *j* possible crossovers occur (in the gamete *i*), *c*_***q***_ is the conditional probability of ***q*** from the Markov model (Figure 4 b) modelling the chromatid interference, i.e.

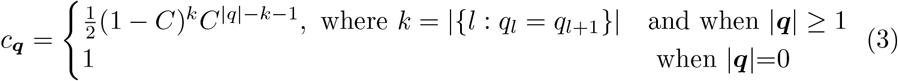

and *C* is the chromatid interference parameter, and *B*_*j*_(*r*_*i*_, ***q***) models the crossover locations, independently for each *j*. This is clearly a generalisation of (1) as by setting (*C* = 0.5 or) *c*_***q***_ = 2^−|***q***|^ and *B*_*j*_(*r*_*i*_, ***q***) = 1(***q*** contains exactly |*r*_*i*_| ones), we obtain the same likelihood function as in (1).

As the model is based on likelihoods, it enables likelihood ratio tests, e.g. for crossover interference (null hypothesis: *B*_*j*_ does not depend on *j*), obligatory crossover (null hypothesis: *p*_0_ = 0) and chromatid interference (null hypothesis: *C* = 0.5).

#### 2.3.1 Non-uniform crossover location distribution and crossover interference

The crossovers locations in the physical coordinates, term *B*_*j*_(*r*_*i*_, ***q***), are modelled as a piecewise constant (relative) rates ***h*** = (*h*_1_, *h*_2_, …, *h*_|***h***|_), *h*_*k*_ ≥ 0, ∑_*k*_ *h*_*k*_ = 1 over an user-defined number of equal width bins (parameter numBins = |***h***|) over the chromosome length. The position distribution of exactly one tetrad crossover is given by ***h***, i.e. 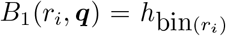, where |***q***| = 1, *q*_1_ = 1 and bin(*r*_*i*_) ∈ {1, 2, …, |***h***|} is the bin of the crossover position *r*_*i*_ (|*r*_*i*_| = 1).

When *j* > 1 crossovers occur, we define a preferred location *m*_*k*_ for each crossover *k*, 1 ≤ *k* ≤ *j*. Instead of using probabilities ***h***, we use a separate ***h***_*k*_ for each crossover *k*. ***h***_*k*_ is obtained by multiplying (and then normalising) ***h*** with a Gaussian density with mean *m*_*k*_ and a standard deviation of 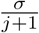. The means *m*_*k*_ (1 ≤ *k* ≤ *j*) are defined as

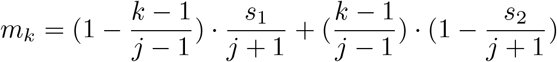

where *s*_1_ and *s*_2_ control the equal spacing of the crossovers. The *s*_1_ controls how close to the start of the chromosome crossovers span and *s*_2_ similarly to the chromosome end, see Figure 3. We also add value *b* to the binned Gaussian densities to allow more variation to the crossover locations than by Gaussian density only. We use dynamic programming to calculate the exact distribution for each of the crossovers, e.g. the first crossover of a total of three. This is necessary with chromatid interference (*C* ≠ 0.5) as *c*_*q*_ varies with same *q* and the different patterns | *q* | do not cancel out in the likelihood. In this way, we define piecewise constant rates for each of the *j* crossovers with crossover interference.

**Figure 3:**
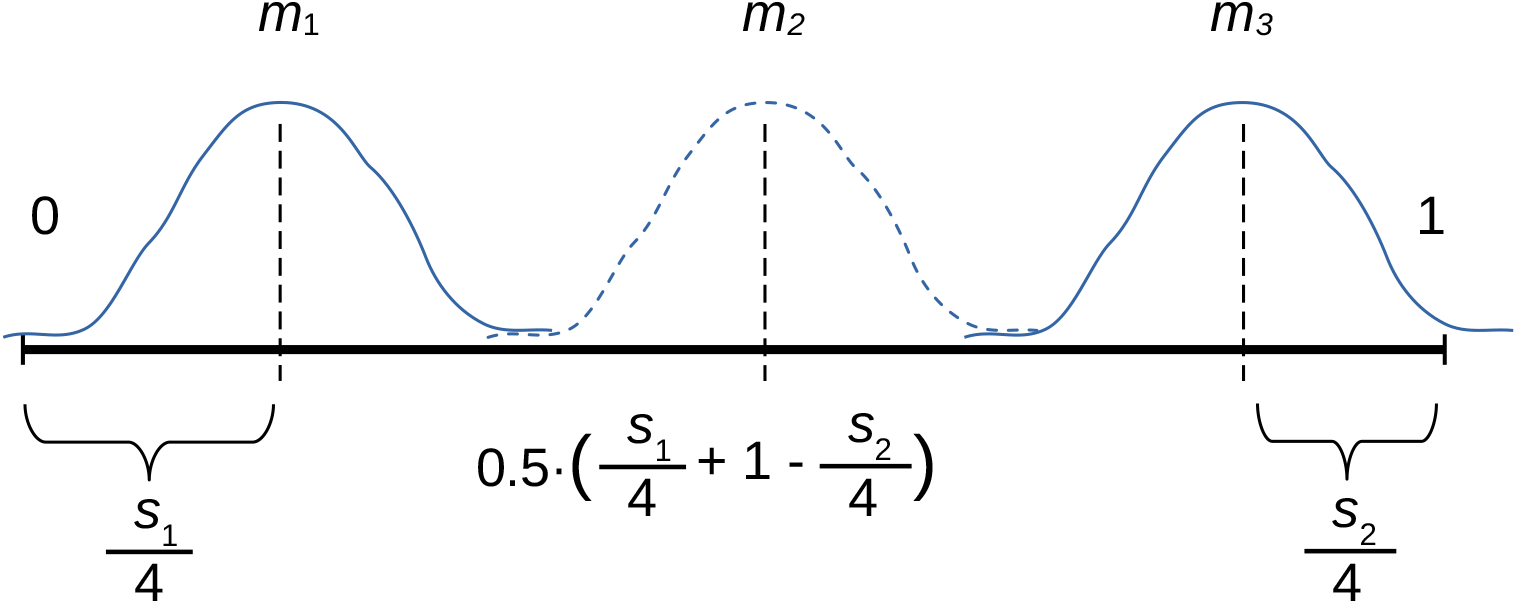
The placement of *j* = 3 crossovers in Lep-Rec. The parameters *s*_1_ and *s*_2_ control how close to the chromosome ends the first and last crossover preferably occur. The remaining *j* − 2 crossover(s) are preferred to occur equispaced between first and last preferred crossover locations. Gaussian kernels with standard deviation of 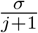 are placed at the preferred positions from 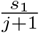 to 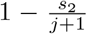 and the variable recombination rate is multiplied by each of these kernels.

Then the crossover position(s) *r*_*i*_ are binned to 1 … | ***h*** | and their likelihoods can be calculated from the corresponding rates indicated by the crossover pattern ***q*** (value of *q*_*l*_ = 1 indicates that the crossover *l* was sampled to the gamete of individual *i*). As final extension, there can also be *H* (parameter numHistograms; default = 1) independent rates ***h*** for exactly 1 … (*H* − 1) and ≥ *H* crossovers. The free parameters ***h***, *σ, b, s*_1_ and *s*_2_ are learned by maximum likelihood method.

This crossover interference model was inspired by the butterfly data, see Figure 6. Its main difference to the existing models (McPeek and Speed, 1995; Otto and Payseur, 2019) is that we model the crossovers in the physical coordinates and do not directly model the crossover distances, only the preferred locations of the crossovers.

We measure the strength of crossover interference concretely as the median of the physical distance *R* of two crossovers from our model. This distance can be contrasted to the median of the distance *R*_0_ of two crossovers sampled directly from the histogram ***h***. We propose the ratio of these median distances as interference score, describing how much the crossover distance increases due to interference.

Another option would be to take the ratio of variances of unit normalised distances (mean = 1) of *R*_0_ and *R*; We denote this ratio as *ν*_Lep-Rec_. This ratio is related to the parameter *ν* in the gamma model as the crossover distances distributed as gamma(1, *λ*) and gamma(*ν, λ · ν*) have variances 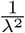 and 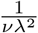, respectively, and hence their ratio is *ν*.

Possible map errors are modelled by deleting or inserting crossovers into the input. This is controlled by two parameters, *ϵ*_1_ for deletion of a marker and *ϵ*_2_ for insertion of a stretch of markers creating either one or two additional crossovers. These parameters multiply the likelihood correspondingly, and the deletion of either one or two crossovers affects the likelihood as many times as there are supporting markers for these crossover(s). The error parameters are given by the user, defaults are 0.1 and 0.001, respectively (deletion of three markers has the same likelihood contribution as one insertion).

#### 2.3.2 Combining our model with gamma sprinkling

It is possible to plug-in the gamma sprinkling model (Copenhaver et al., 2002; Housworth and Stahl, 2003) into formula 2, as the gamma sprinkling model naturally defines the probabilities *p*_*i*_ and *B*_*j*_(*r*_*i*_, ***q***). We have implemented these computations by discretising the relative chromosomal region [0, 1] into (numBins=30) bins and calculating the required probabilities via dynamic programming. As simplification, we assume here that the crossover distance is a mixture of gamma and exponential distributions, not that a fraction *p* of crossovers would be sampled according to the Poisson process (exponential distance) and the remaining according to the gamma model disregarding the Poisson distributed crossovers. Moreover, it would be possible to learn parameters *p*_*i*_ freely, but define *B*_*j*_(*r*_*i*_, ***q***) with the gamma sprinkling model, or vice versa. We also obtain the gamma model (McPeek and Speed, 1995) by restricting the mixture fraction *p* = 0. We call these simplified gamma and gamma sprinkling models as these do not directly model the crossover distances (correlations) of adjacent crossovers. We have also implemented full gamma and gamma sprinkling models (with discretised crossover distances and hence correlations for adjacent crossovers) for our experimental comparisons.

#### 2.3.3 Chromatid interference

The chromatid interference is modelled by a parameter *C* in Lep-Rec. This parameter is the transition parameter in a Markov model explained in Figure 4 (b). The value *C* = 0.5 means no interference, whereas *C <* 0.5 indicates negative and *C* > 0.5 positive interference. The relation between *C* and the probability *c*_***q***_ of gamete’s crossover pattern ***q*** (conditional on | ***q*** |) is defined by the Equation 3.

**Figure 4:**
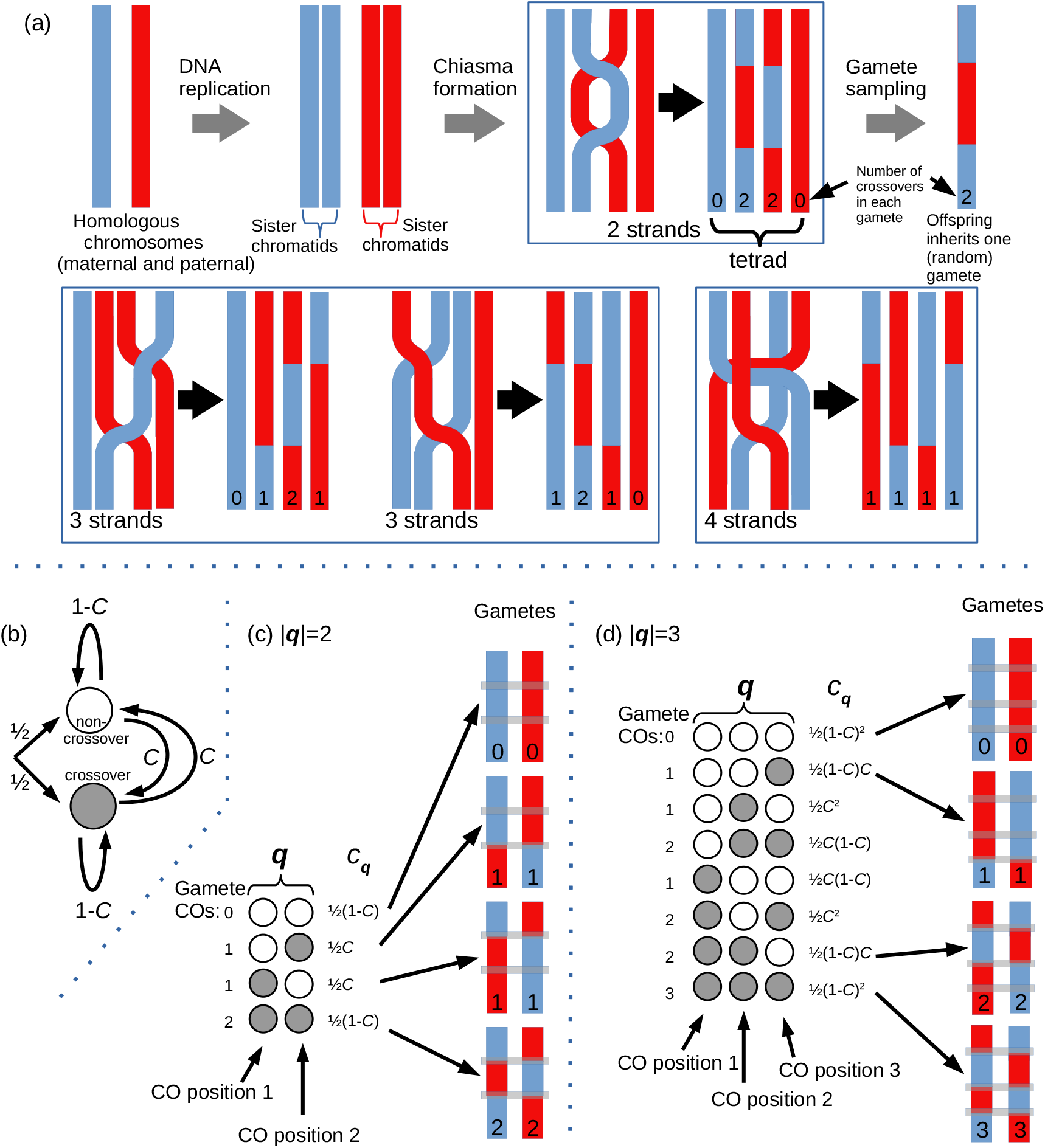
(a) A simplified diagram of meiotic recombination and the four possible outcomes from tetrad double crossover, inspired by Sarens et al. (2021). Chromatid interference (CI) affects how the sister chromatids participate in the crossovers favouring either 2-strand (same chromatid pair in both crossovers) or 4-strand cases (disjoint chromatid pairs). Without CI, the four outcomes should be equal likely. (b) The Markov model used to model CI in Lep-Rec, the grey states corresponding to sampled crossovers (and to values of 1 in ***q*** of Equation 2). The parameter *C* controls CI, when *C* = 0.5 there is no CI, *C* > 0.5 indicates positive and *C <* 0.5 negative interference. (c) The conditional probabilities of different offspring gametes originating from two tetrad crossovers. (d) Similar figure as (c) showing only some gametes for three tetrad crossovers.

#### 2.3.4 Learning the model

All parts explained above have been implemented into a single model and maximum likelihood (ML) method is used to fit the free parameters *p*_*j*_ for the rate of tetrad crossovers, *C* for chromatid interference, *b, s*_1_, *s*_2_ and *σ* for crossover interference and the binned probabilities ***h*** for variable recombination rate. Parameters *p*_*j*_ and *C* are learned in a typical way using an EM algorithm. However, the remaining parameters are more difficult; in order to optimise these, each of them is increased or decreased separately and parameter value is changed if the expected likelihood is improving. In the case of the gamma sprinkling model, parameter *C* is learned by the EM algorithm, whereas the parameters *ν, λ* and *p* are learned as these remaining parameters in our model.

The runtime for one EM iteration is 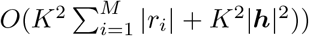, where individual *i* has *r*_*i*_ crossovers and the learned histogram ***h*** has |***h***| *>* 1 bins. For simplified gamma and sprinkling models, the runtime is slightly faster 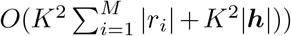. To learn these parameters, Lep-Rec runs about 1600 iterations from multiple initial parameter values, out of which 1000 iterations contribute to the final result (controlled by the parameter maxIterations=1000).

### 2.4 Data

In order to evaluate Lep-Rec performance on real data, we constructed linkage maps for two distant animal groups, butterfly and fish. We collected crosses that could be included in the maps from several whole-genome and reduced representation sequencing (RAD-seq, DArTseq) studies in order to maximise the number of observed crossovers. The corresponding sequencing reads were retrieved from public repositories (Supplementary tables 2 and 3) and mapped to the reference genomes using the program BWA mem (Li, 2013).

The fish map was constructed by combining data from two nine-spined stickleback (*Pungitius pungitius*) studies (see Supplementary table 2). The sequencing data was mapped to the NSP V7 (accession GCA 902500615.3) genome assembly.

The butterfly map was constructed by combining data from two *Heliconius* species across several studies (see Supplementary table 3). The two species, postman butterfly (*Heliconius melpomene*) and longwing cydno (*Heliconius cydno*) have extremely similar genome structures and recombination landscapes (Davey et al. (2017)). Thus, all the reads were mapped to the postman butterfly genome (Davey et al. (2016), version Hmel2.5).

Both variant calling and map construction were done using Lep-MAP3 modules (Rastas, 2017). The variant calling used the mpileup command of the samtools program (version 1.13) together with the Lep-MAP3 module Pileup2Likelihoods as described in the Lep-MAP3 documentation. Next, the IBD module was used to infer the pedigrees and the results were checked against available annotations. Individuals and families were removed if their relationships remained uncertain. The final fish data set contained 1,695 and butterfly data set 1,007 offspring, respectively.

Then the module ParentCall2 was used to call the parent genotypes. The fish parent genotypes were called with option Xlimit=2 and butterfly with Zlimit=2 to find X and Z inherited markers in accordance with the respective inheritance systems of the species. Next, the module Filtering2 was used to filter problematic variants. Additionally, if consecutive physical positions in the reference genome had variants, only the first one was kept. After running the module SeparateChromosomes2 with a range of logarithm of the odds (LOD) score limits, the results were manually checked and the smallest LOD scores were chosen that divided the genome into major linkage groups that matched the 21 reference chromosomes (1n=21). Additional markers were added to the fish linkage groups using the module JoinSingles2All.

The markers within linkage groups were ordered by running the module OrderMarkers2 three times for each linkage group and taking the order that had the highest likelihood. The effect of possible genotyping errors originating from difficult genome regions was mitigated by scaling markers within 100 bp from each other so that their combined effect on linkage map was approximately the same as one marker without any other markers within the same distance(option proximityScale=100). The resulting *de novo* linkage maps were post-processed using the module FitStepFunction. Additional linkage maps were generated by evaluating the markers in each linkage group in the physical order determined by the reference genome by rerunning the OrderMarkers2 module. Butterfly markers were ordered with option recombination2=0 in accordance with absence of female recombination in the species. The resulting fish and butterfly linkage maps contained 91,581 and 211,515 markers, respectively.

Additionally, we used a previously published data on cattle (*Bos taurus*) with individual crossover locations over the autosomes (Kadri et al., 2016). This data contains 25332 maternal (dam) and 94516 paternal (sire) crossover locations (in Mb) for 29 autosomes. We used a custom script to convert these crossover locations to the output format of OrderMarkers2 with 4 markers per each Mb (chromosome lengths were 43-158 Mb, mean=86 Mb).

The centromeres separate these species, as butterflies do not have centromeres (Suomalainen, 1966), whereas fish and cattle do have them. Moreover, the centromere location is variable only in the fish as all cattle autosomes are acrocentric (De Lorenzi et al., 2017). As final difference, the cattle data is significantly larger than the two other and allows association analysis for the recombination rate.

We also simulated data to evaluate how our method can estimate crossover patterns, crossover and chromatid interference and tetrad crossover distributions. Especially, we compared our method against the method of Yu and Feingold (2001) (and its generalisations) which ignore the crossover locations.

## 3 Results

First, we evaluated FitStepFunction by comparing the recombination rates obtained from maps after FitStepFunction and the physical maps. The both ways obtained comparable results, Figure 5 contains the two rates for fish chromosome 4 and two spline fits for a comparison. With the spline fits, we obtained negative recombination rates, even when using maps consistent with the physical order (by FitStepFunction or the physical maps).

**Figure 5:**
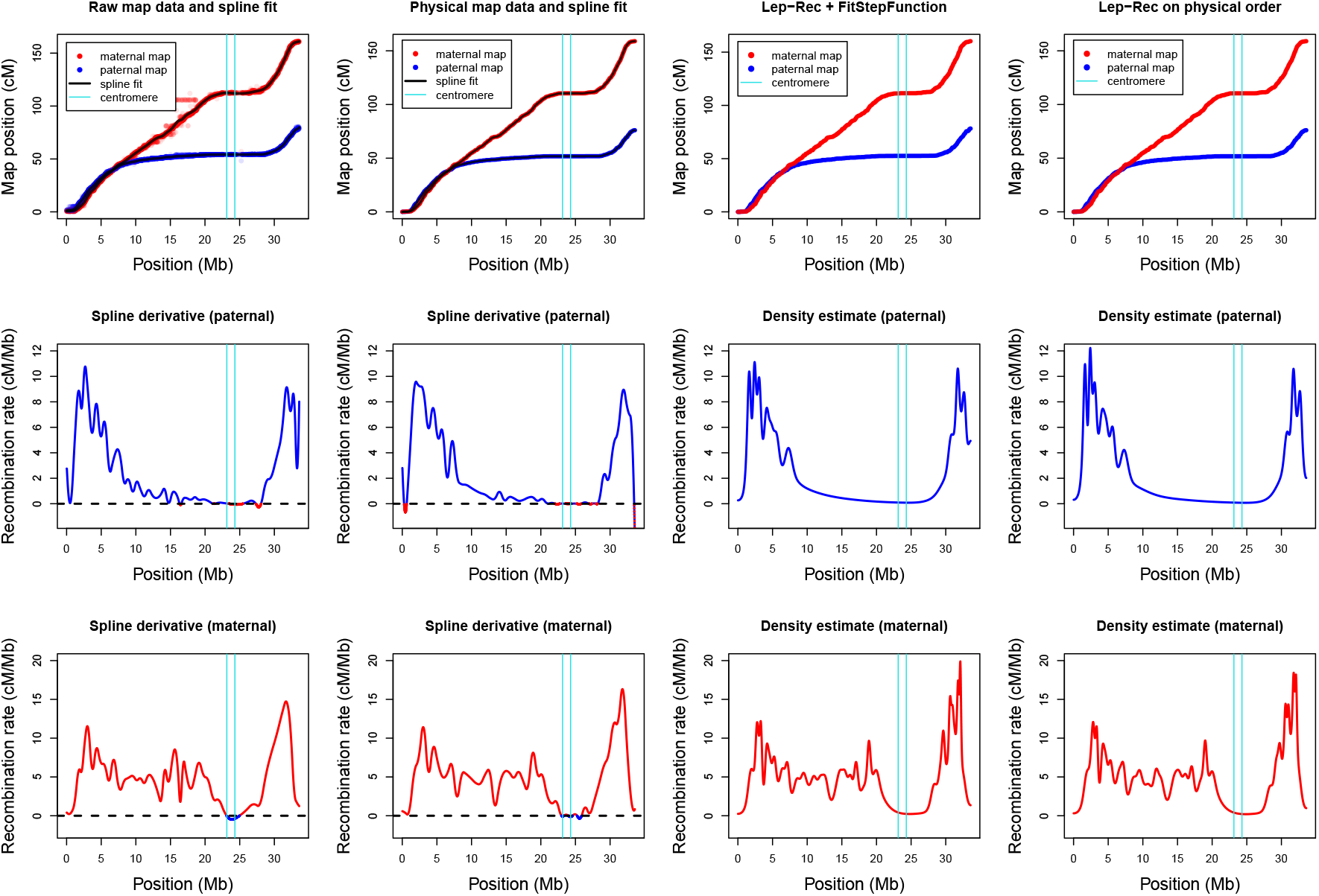
Estimated recombination rates for nine-spined stickleback’s longest chromosome 4. The left column displays the raw map data, with spline fits and their derivative as the recombination rate estimates, whereas the second column has the same estimates using the map in the physical order. The third column contains results with FitStepFunction from the raw map (and map intervals) and the recombination (density) estimates obtained by Lep-Rec’s EstimateRecombination module. The rightmost column shows the maps evaluated in the physical order and their recombination estimates by EstimateRecombination. Smoothness parameter *S* = 40 was used for density estimation (parameter bandwidth=-40) in order to obtain a similar smoothness as the spline fit (cross-validation yielded much smaller *S* and more spiky fit). The centromere region from Kivikoski et al. (2021a) is indicated by the cyan lines.

The physical order was used in our tetrad analysis. We learned our tetrad model for each data set with 8 parameter combinations enabling or disabling crossover and chromatid interference and obligatory crossover (*p*_0_ = 0 or *p*_0_ *>* 0). The butterfly data was analysed with and without the error model in Lep-Rec, all other tetrad analysis was conducted only with the error model enabled.

We also learned simplified and full gamma and gamma sprinkling models for each data set with the same parameter combinations. Crossover interference was disabled by setting parameter *ν* = 1 (Poisson process), and obligatory crossover was enabled by conditioning the model for *p*_0_ = 0 (dividing each *p*_*i*_ by 1 − *p*_0_ and then setting *p*_0_ = 0). In each case, we used the data likelihood to compare different models. To decide between the parameter combinations, we used likelihood ratio test (chi-square distribution with *k* free parameters, twice the log likelihood difference 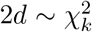). We used p-value *<* 0.001 for each comparison, leading to likelihood differences of 6.6 and 3.3 (natural logarithm) for degrees of freedom of *k* = 4 (Lep-Rec’s crossover interference with 4 parameters) and *k* = 1 (single parameter, like *ν* ≠ 1), respectively. We plotted these likelihoods (and some fitted parameters) in the descending order of the chromosome length, which is a reasonable proxy for the map length (e.g. Rastas (2017)) and also for the strength of crossover interference (Otto and Payseur, 2019). Moreover, longer map has more crossovers making the data more informative. Separate results were obtained for male and female parents, except for butterfly, in which only the male parent recombines (Suomalainen, 1966).

### 3.1 Butterfly

Our crossover interference model was inspired by the butterfly data. This was ideal data for interference study as the recombination rate was relatively constant across chromosomes and the map lengths were relative short (47-66 cM, average 55.5 cM). Moreover, butterflies do not have centromeres, so we can ignore their (decreasing) effect on recombination.

For the longest chromosomes (≈60 cM) we could clearly see the effect of crossover interference as two “bumps” in the recombination rate at relative positions of about 1/5 and 4/5. As an example, Figure 6 has data from chromosome 10 with the longest map of about 66 cM. From this example, we could also estimate the distance of crossovers from the 71 gametes with exactly two crossovers, since these are most likely from two tetrad crossovers (there were no individuals with three or more crossovers). To our surprise, the linear correlation (about 0.23) between the first and second crossover positions (in Mb, similar result in cM) was not significant (*p* = 0.0542)! However, the distributions of the first and second crossover have hardly any overlap (Figure 6 (c)), suggesting strong crossover interference. For the other 8 chromosomes with at least 30 double crossovers, only two chromosomes (6 and 19) had significant correlations (p=0.013 and p=0.0052). The combined p-value for these correlations was significant (p=0.007, Fisher’s method (Fisher, 1925)), but clearly the correlation was fairly weak, as only two of the 9 correlations were significant. This supported our simple crossover interference model: The distribution of crossover positions and their distances are bell-shaped and crossover positions are not highly correlated. However, there can be interference without (linear) correlation.

**Figure 6:**
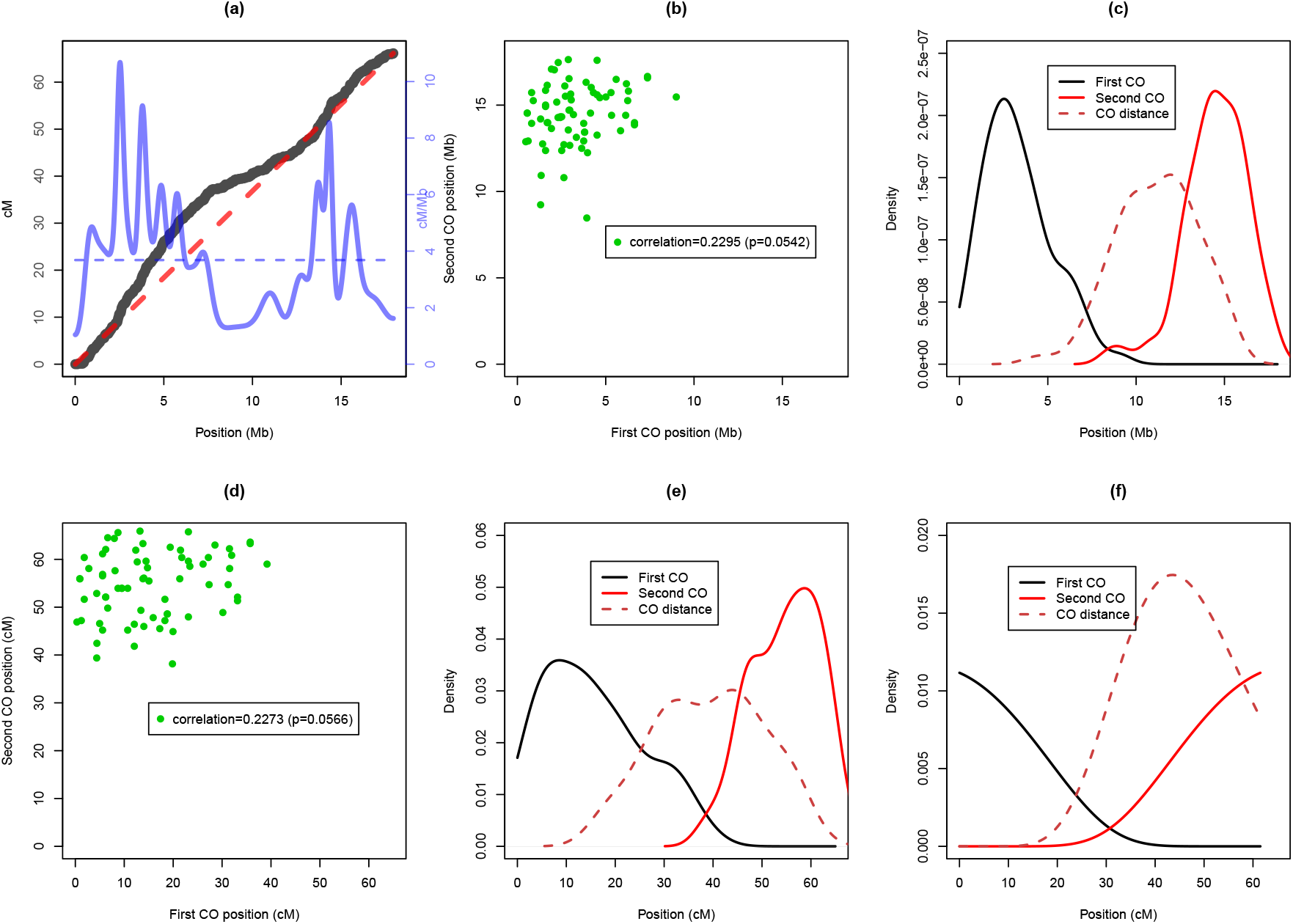
Butterfly data for chromosome 10. Panel (a) displays the corresponding Marey map (black) and recombination rate (blue). Dashed blue line shows the average recombination rate, which as constant recombination rate would correspond to the dashed red line as Marey map. Panel (b) plots crossovers positions in gametes with exactly two crossovers. The densities of these positions and their distance are shown in panel (c). Panels (d) and (e) show the same crossover positions in cM. (f) The fitted gamma (sprinkling) model (*λ* = 1.2958 *ν* = 12.191, *p* = 0.0).

Figure 6 (f) shows the crossover distributions from a maximum likelihood fit of the gamma sprinkling model (simplified and full model gave almost identical parameters). These distributions are not very similar to the actual two crossover data. Moreover, simulating data from the fitted gamma sprinkling model generated higher (*>*0.4) and more significant correlations for the first and last crossover (of two) as observed in the real data.

Figure 7 has the results for the tetrad analysis. Enabling crossover interference improved Lep-Rec’s likelihood much more on the 11 longest chromosomes than on the 10 shorter ones. Most of these likelihood differences were still significant, except for chromosomes 15, 16 and 4. However, enabling *p*_0_ *>* 0 did not significantly improve the likelihood. There were only two chromosomes (18 and 6) where allowing chromatid interference significantly improved the likelihood, in both cases the parameter *C* was close to zero. Without the error model, chromosomes 16 and 19 also have significant likelihood improvements with *C* ≠ 0.5. The conclusion of our analysis is that there is evidence for crossover interference, but no evidence for non-obligatory crossover (*p*_0_ *>* 0). The chromatid interference might have a small effect on some chromosomes, but we cannot be absolutely sure as the map errors seem to cause similar signals in the data with our analysis. Moreover, extremely strong negative interference (*C* ≈ 0) seems unlikely.

**Figure 7:**
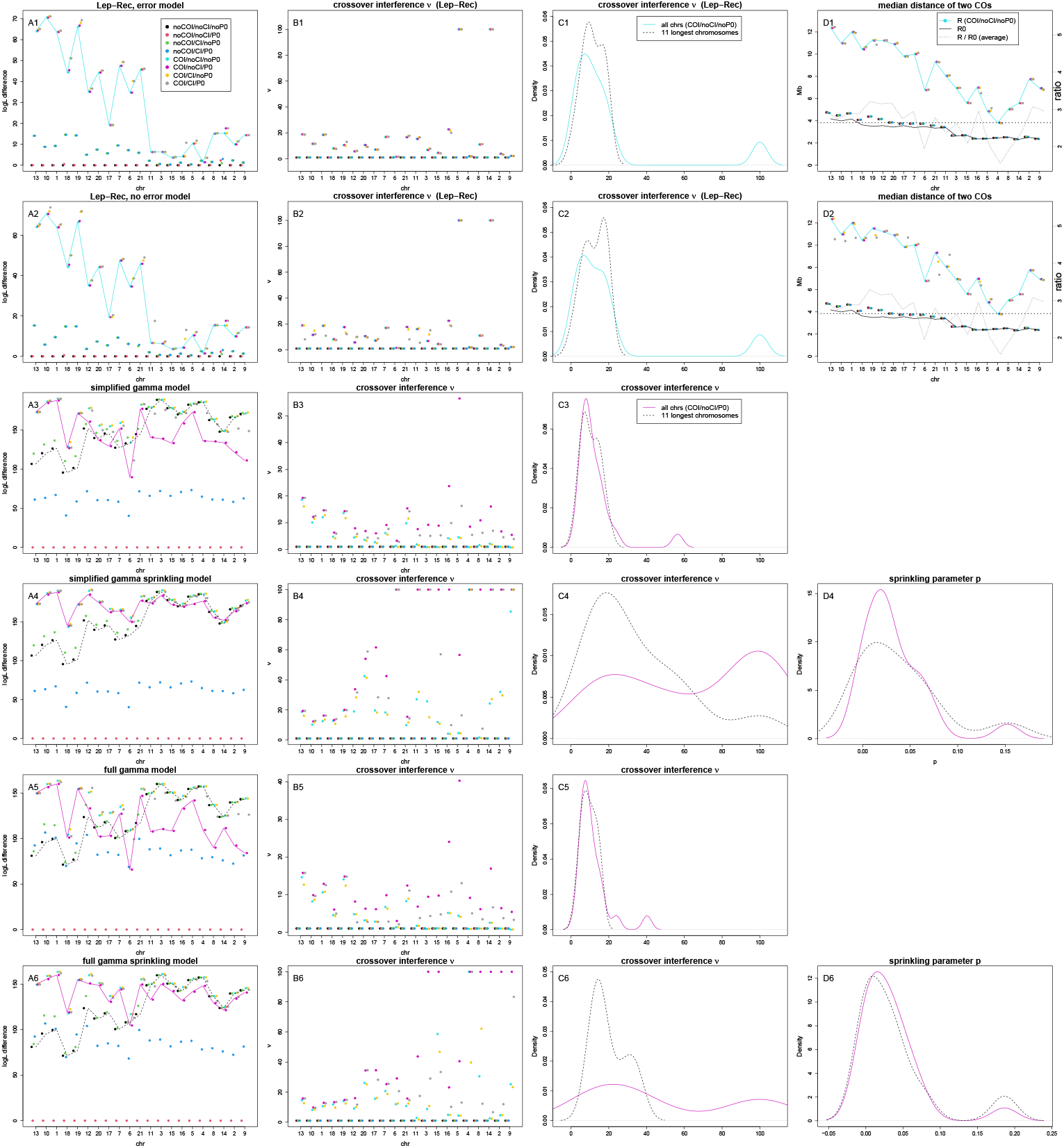
Tetrad analysis on butterfly data. First two rows use Lep-Rec’s model with and without error modelling. Third and fourth row are using simplified gamma and gamma sprinkling models and the last two full gamma (sprinkling) models, all with error modelling. Panels A1-A6 display the likelihood differences (support) for each 8 combinations of enabling/disabling crossover (COI) and chromatid interference (CI), and zero crossover tetrads (P0). Panels B1-B6 have the estimates for crossover interference parameter *ν* for each chr, whereas C1-C6 depicts the density of the estimated *ν* parameters over all chromosomes with COI/noCI/noP0 (Lep-Rec) and COI/noCI/P0 (gamma and gamma sprinkling). The panels D1 and D2 contain the median distances of two crossovers in Lep-Rec’s models, and the densities of estimated parameters *p* in the gamma sprinkling models are in D4 and D6. Chromosomes are in descending length-wise order.

With the gamma and sprinkling models, the likelihood difference between *ν* = 1 and *ν* > 1 was negatively correlated with chromosome length, indicating that the model is not very sensitive to the crossover distances. This could be because the data contains too few crossovers to learn these models or because the crossover interference in these models also accounts for the obligatory crossover (*p*_0_ = 0). Conditioning these models on *p*_0_ = 0 improved the likelihood on almost all chromosomes. Possibly, such conditioned (Poisson) model provides a better contrast for the likelihoods (separating the effect of interference between two or more crossovers and obligatory crossover) and is also plotted as dashed line to the Figure 7. Chromatid interference was significant on several chromosomes, when comparing likelihood difference against normal gamma (sprinkling) models with or without CI. However, when comparing against models with obligatory crossover, only the same chromosomes as before, 18 and 6, remained with significant likelihood improvement. Moreover, the full gamma (sprinkling) model improved with chromatid interference with *ν* = 1 (Poisson). This was because strong negative interference (*C* ≈ 0) selects every second crossover and effectively obtains a better fitting gamma distance for crossovers with *ν* = 2.

The difference between gamma and gamma sprinkling models was that the likelihood of the sprinkling model was not improved much when conditioned with *p*_0_ = 0, e.g. it can better model obligatory crossover by increasing *ν* and *p*. This can be seen in Figure 7 as its likelihood follows closely to the conditioned Poisson model for the shortest chromosomes.

Moreover, it was difficult to decide a value for *ν* due to the high variance of the parameter estimates between chromosomes and between the gamma and gamma sprinkling models (parameter values for *ν* are reported for many species by e.g. Otto and Payseur (2019)). The estimates of *ν*_Lep-Rec_ and *ν* of the gamma model were similar, both varied less on different chromosomes than the gamma sprinkling model. The highest estimates of *ν* were discarded by considering only the 11 longest chromosomes. The ratio of the median distances of *R*_0_ and *R* was on average 2.6, i.e. the crossover distance increased by 160% with crossover interference.

### 3.2 Fish

The fish female map lengths were 77-159 cM (average 104 cM), male maps 51-76 cM (average 56.5 cM). As depicted in Figure 5, the fish centromere location is easily seen from the Lep-Rec’s recombination rate as the rate drops to near zero in this region while remaining non-negative. On the other hand, the recombination rate from a spline fit has more fluctuations and negative values in these regions. Moreover, spline fit yields negative values also at the map ends.

The figures 9 and 10 have the results of the tetrad analysis. For female maps, the likelihood difference for crossover interference (*ν* = 1 and *ν* > 1, or Lep-Rec’s 4 parameter model) was correlated with the chromosome length in all models. All chromosomes and models show significant crossover interference. The estimates of *ν* were smallest in the gamma and largest in the gamma sprinkling model. The ratio of median distances of *R*_0_ and *R* was on average 3.0, i.e. the crossover distance increased by 200% with crossover interference.

Lep-Rec’s model did not support *p*_0_ *>* 0, but chromatid interference was significant on three chromosomes, chr1 (*C* = 0.58), chr2 (*C* = 0.55) and chr18 (*C* = 0.41). The likelihoods of the gamma model improved generally with *p*_0_ = 0, especially on the shortest chromosomes. The simplified gamma model obtained significant chromatid interference on chromosomes 16 and 18, the simplified gamma sprinkling on 13 and 16. To our surprise, the full gamma and gamma sprinkling models found many more chromosomes with significant chromatid interference but not on the same chromosomes as on the other models.

For male maps, Lep-Rec’s model on crossover interference obtained significant likelihood differences on all chromosomes, but the differences were much larger on about half of the chromosomes without a clear pattern on the chromosome length. Chromatid interference improved the likelihood on several chromosomes, but there was no support for non-obligatory crossovers (*p*_0_ *>* 0).

The gamma model’s likelihood difference between *ν* = 1 and *ν* > 1 was positively correlated with the chromosome length, for the gamma sprinkling there was no clear length correlation at all. The likelihoods of the gamma model improved by conditioning with *p*_0_ = 0. Especially the sprinkling model obtained highly variable estimates for *ν*, often obtaining the maximum value of 100 in our implementation. The estimates of Lep-Rec and the gamma model were more similar, but gamma model obtained smaller estimates of *ν*. The ratio of median distances of *R*_0_ and *R* was on average 4.9, i.e. crossover distance increased by almost 400% with crossover interference (much higher compared to 200% in female, possibly explaining the shorter male map).

Simplified gamma and gamma sprinkling models supported chromatid interference on several chromosomes. To our surprise, and opposite to the female maps, the full gamma and sprinkling models did not support chromatid interference (except sprinkling model for chr16). This variable support from different models could be due to how well each model fits to the data. Moreover, these short maps do not present strong evidence for chromatid interference as it involves two or more tetrad crossovers. Note that the outlier chr12 is the sex chromosome with a long non-recombining region in the male map.

### 3.3 Cattle

The cattle dam map lengths were 50-123 cM (average 78.9 cM), sire maps 52-128 cM (average 87.3 cM). As there was plenty of data, we increased Lep-Rec’s parameters to *K* = 10 and |*h*| = 30 (maxCrossovers=10, numbins=30, defaults are 8 and 10) in the tetrad analysis. The likelihood difference was significant for crossover interference on every chromosome, and the difference was correlated with the chromosome length in all models and for sire and dam. In Lep-Rec and the gamma model, the estimates for *ν* were positively correlated with the chromosome length as well. For the full gamma sprinkling model, the correlation was negative as reported by Otto and Payseur (2019), but the estimates of the 4 smallest chromosomes might be outliers. However, this correlation is not as apparent for the simplified gamma sprinkling model. Generally, there was no clear correlation between the parameter *p* and the chromosome length. The average ratios of the medians of distances *R*_0_ and *R* were 3.3 and 3.0, for dam and sire, respectively (also higher in dam, where the map length is shorter). Moreover, the ratios were positively correlated with the chromosome length.

The difference in the Lep-Rec’s likelihoods suggested that crossovers might be non-obligatory for some of the shortest chromosomes (20, 23-29). For these same chromosomes, the chromatid interference was also highly supported. However, the shortest chromosomes should be the least informative for chromatid interference (requires two or more tetrad crossovers) so we do not take this as evidence of this, but probably related to non-obligatory or missing crossovers in the data. Few longer chromosomes showed evidence for chromatid interference as well. However, using the gamma and sprinkling models, almost every chromosome supported chromatid interference, without difference between simplified and full models.

We also calculated the mean number of tetrad crossovers for each offspring using Lep-Rec without chromatid interference nor non-obligatory crossover. We call this measure as (mean) tetrad recombination rate (TRR) and compared it to the sum of gametic crossovers (global recombination rate, GRR) used by Kadri et al. (2016). The density of (gamete) crossovers and tetrad crossovers is shown in Figure 13, TRR having a standard deviation of about half of the deviation of GRR. Then we recalculated the association with the parental genotype data for both GRR and TRR. Table 1 shows that TRR obtains more significant association with the genotypes, especially for sire. We also calculated the heritability of these recombination rates from a simple linear model

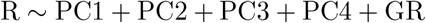

where R and GR were the parent’s and its grandparent’s (dam of dam/sire of sire) recombination rate (GRR or TRR). The variables PC1 to PC4 are the principal components describing the population structure. The heritability was taken as twice the regression coefficient of GR. As Kadri et al. (2016), we took only a random trio from each family. This way, we obtained heritabilities of 0.182 and 0.156 for TRR, for sire and dam, respectively. For GRR, the same values were 0.140 and 0.0914, for sire and dam, respectively, being comparable to the values reported by Kadri et al. (2016) (0.13 *±* 0.03 and 0.08 *±*0.02). Thus, heritablilities for TRR were 30% and 70% higher than for GRR.

**Table 1:** Association of SNPs and tetrad recombination rate (TRR) and global recombination (GRR) on the data of Kadri et al. (2016). Each cell gives the coefficient of the linear model ([TG]RR∼ PC1+PC2+PC3+PC4+SNP1+… +SNP11) and the corresponding p-value. The lowest p-value is in **bold**. *=based on a sample of 50% of the data

| SNP | sire TRR | sire GRR | dam TRR | dam GRR |
| --- | --- | --- | --- | --- |
| HFM_S1189L | -0.279/**3.42e-084** | -0.581/2.98e-081 | -0.346/**8.63e-23** | -0.608/7.39e-22 |
| MSH4_lead | -0.010/**5.64e-001** | 0.007/8.45e-001 | -0.300/4.15e-11 | -0.567/**4.14e-12** |
| MSH4_C342Y | -0.180/**7.04e-033** | -0.306/1.10e-021 | -0.057/**1.94e-01** | -0.089/2.58e-01 |
| RNF212_P259S | 0.457/**1.22e-252** | 0.925/9.34e-231 | 0.352/9.64e-25 | 0.644/**1.30e-25** |
| RNF212_A77T | -1.245/**2.43e-204** | -2.340/4.71e-161 | -0.291/**6.83e-02** | -0.478/9.58e-02 |
| RNF212B_1 | 0.742/**0.00\*** | 1.423/0.00 | 0.565/**4.33e-39** | 0.955/7.57e-35 |
| RNF212B_M2 | -0.358/**5.47e-217** | -0.713/2.39e-191 | -0.083/5.13e-03 | -0.169/**1.47e-03** |
| RNF212B_F2 | -0.109/**5.80e-006** | -0.223/1.31e-005 | -0.485/3.78e-09 | -0.893/**1.51e-09** |
| MLH3_N408S | 0.413/**4.31e-269** | 0.719/5.45e-182 | 0.292/**7.84e-24** | 0.493/2.87e-21 |
| BTA18_rs135... | -0.338/**4.24e-184** | -0.677/2.95e-164 | -0.101/**3.26e-03** | -0.135/2.84e-02 |
| MSH5_R631Q | -0.452/**3.18e-066** | -0.894/6.98e-058 | 0.063/**3.33e-01** | 0.077/5.08e-01 |

### 3.4 Simulated data

The data simulation is explained in Figure 8. We ran Lep-Rec on each simulated data with the default parameters, except varying for numBins, learnCI and leanrCOI as follows. The model of Yu and Feingold (2001) was obtained with Lep-Rec using the parameter “numBins=1 1”, which disables the variable recombination rate (and hence crossover interference). Moreover, chromatid interference parameter *C* was optimised via parameter “learnCI=1” and crossover interference was disabled by “learnCOI=0”.

**Figure 8:**
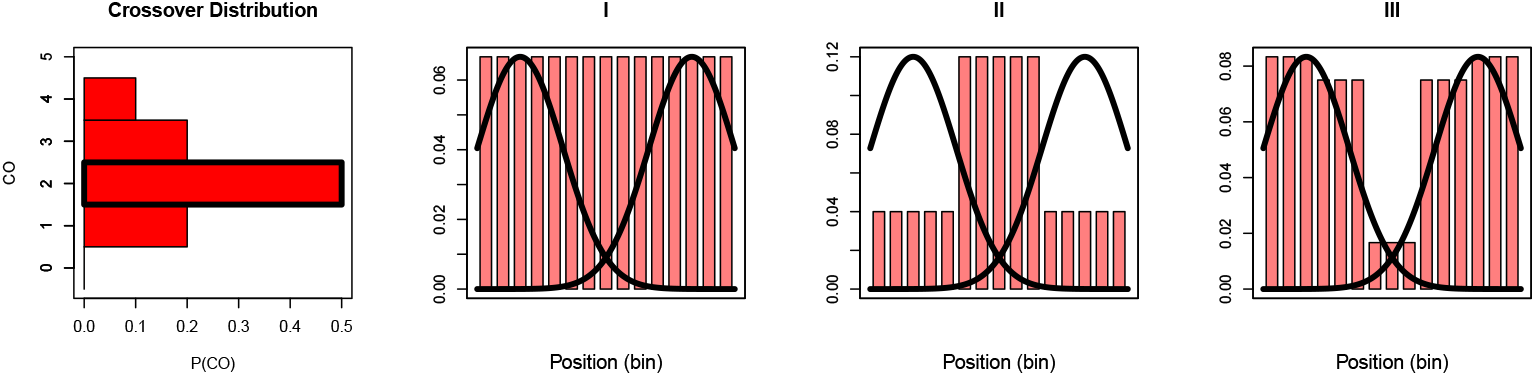
Simulated data. We simulated 1000 offspring and 1000 markers from a single (F1) family to access Lep-Rec’s tetrad analysis performance. The crossover distribution used in the simulations is shown on the left. The other panels (I-III) show the used recombination rates for a single crossover over the chromosome length (as histogram, similar to used butterfly, cattle and fish data, respectively I-III) and crossover interference parameters *s*_1_ = *s*_2_ = 0.5 and *σ* = 0.5 acting on two crossovers (solid black lines). Moreover, the chromatid interference was varied among 0.25, 0.5 and 0.75.

Overall, the variance in the estimated crossover distribution is much smaller when taking into account variable recombination and crossover interference, compared to the model of Yu and Feingold (2001). Moreover, the generalised model of Yu and Feingold (2001) yielded poor estimates of the parameter *C*. Enabling the crossover interference had the largest likelihood improvement in our model, as was the case with the real data.

## 4 Discussion and Conclusion

Our software Lep-Rec provides an easy and objective way to characterise recombination. It is based on the idea of making the physical and linkage positions consistent and then observing the step-wise nature of the data. By utilising this, it obtains robust recombination rate estimates by simple smoothing directly on the crossover locations. This avoids the problems with curve fitting to obtain the Marey map. Lep-Rec’s smoothing is intuitively based either on *S* nearest neighbours or on a fixed physical length. Cross-validation can be used to find parameter *S*, or different values can be tried to obtain the desired smoothness.

We have also shown that Lep-Rec’s tetrad analysis obtains more plausible results on real data (fish, cattle and butterfly) than the widely used gamma and gamma sprinkling models. Our model enables hypothesis testing for crossover and chromatid interference as well as for obligatory crossover (crossover assurance). For example, Lep-Rec’s likelihood difference points clearly when crossover interference has a significant effect in the data, whereas this effect was sometimes difficult to see with the gamma (sprinkling) model. The median crossover distances from Lep-Rec were also easy to interpret, whereas in some of our experiments, gamma and gamma sprinkling models gave opposite correlations with the interference parameter *ν* and chromosome length, as well as sex (differences between male and female maps of the same species). This is probably because only our model separates the effects of crossover interference and obligatory crossover, effects that could be caused by different processes. We have also added chromatid interference and map errors to our model and also implemented these to some of the previously proposed models. Lep-Rec can and should be routinely applied to new and old linkage mapping projects, shedding light on the crossover processes over many species and taxa.

Finally, the estimated crossover distribution can be used to improve heritability estimates and add power to association analysis of recombination. This is possible by obtaining extra information from the crossover locations, and in doing so, reducing the variance caused by (random) sampling of gametes.

For future work, we consider using bootstrapping (Tibshirani and Efron, 1993) to obtain confidence intervals for our model parameters. Another interesting venue would be to output recombination rates according to the Lep-Rec’s model for each number of (tetrad) crossovers. This could be used, e.g. to hypothesise how increased recombination would affect the recombination landscape.

## Acknowledgements

The authors would like to thank Mikko Kivikoski for valuable comments on this manuscript.

---

**Table 2:** Nine-spined stickleback data sets used to construct the linkage map. For each study an identifier, the ENA accession, the references to the original publications, and the number of individuals and families included in the linkage map are shown.

| Study | Accession | References | Ind. | Fam. |
| --- | --- | --- | --- | --- |
| HEL | PRJEB39736,<br>PRJEB39760 | Kivikoski et al. (2021b) | 1216 | 92 |
| HELxPYO/BYN | PRJNA673430 | Kemppainen et al. (2021) | 628 | 2 |
| Total |  |  | 1844 | 94 |

**Table 3:** Heliconius data sets used to generate the linkage map. For each study an identifier, the ENA accession, the references to the original publications, and the number of individuals and families included in the linkage map are shown.

| Study | Accession | References | Ind. | Fam. |
| --- | --- | --- | --- | --- |
| 1 | PRJEB2745 | The Heliconius Genome Consortium (2012) | 48 | 1 |
| 2 | PRJEB11288 | Davey et al. (2016) | 73 | 1 |
| 3 | PRJEB16767 | Davey et al. (2017) | 424 | 4 |
| 4 | PRJEB34160 | Byers et al. (2020),<br>Byers et al. (2021),<br>Darragh et al. (2021) | 337 | 25 |
| 5 | PRJEB38330 | Bainbridge et al. (2020),<br>Brien et al. (2022) | 215 | 3 |
| Total |  |  | 1104 | 34 |

**Figure 9:**
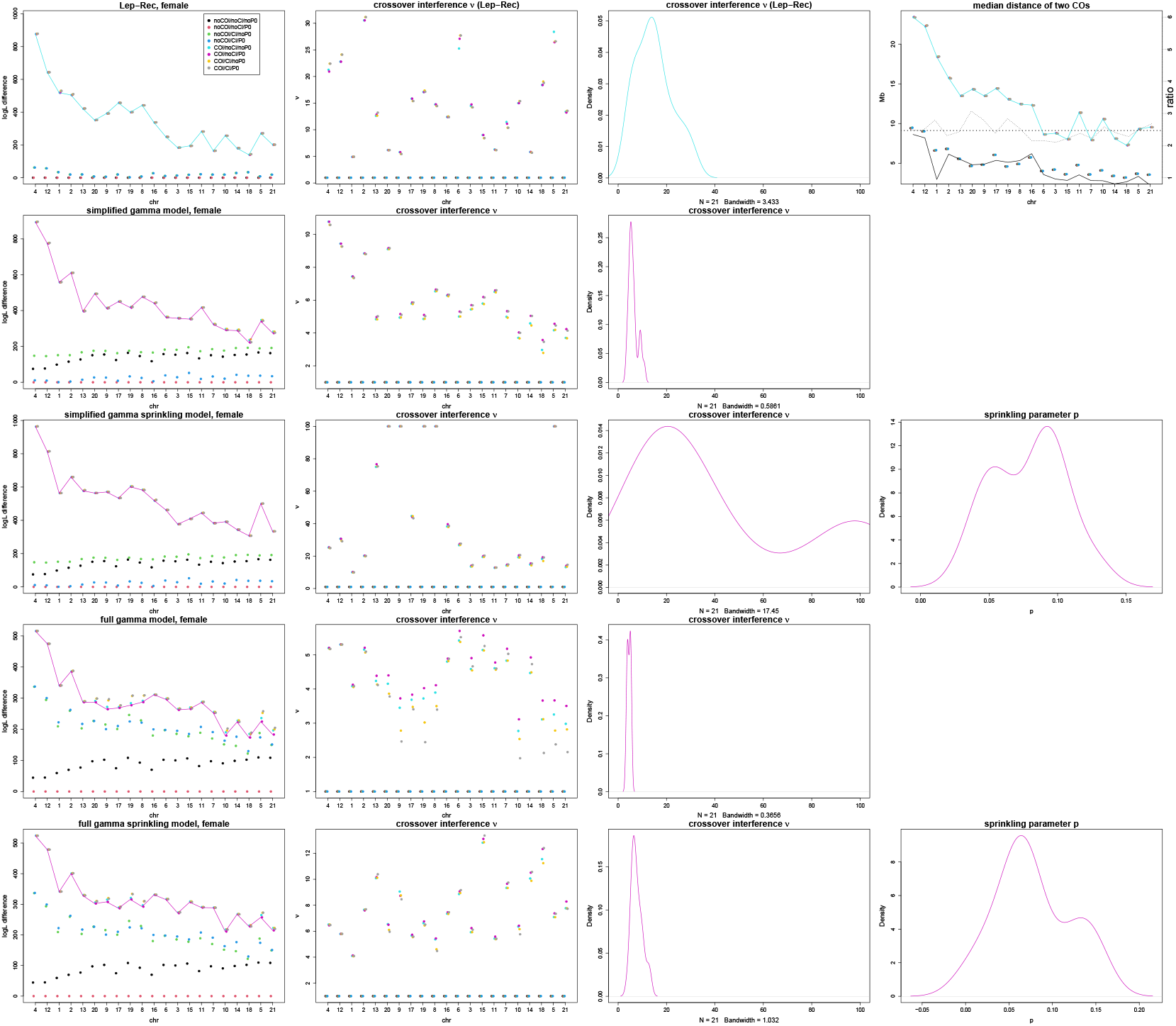
Tetrad analysis on female fish data. The results on the first row are using Lep-Rec’s model, second and fourth row simplified and full gamma models and the third and last row are using simplified and full gamma sprinkling, all with error model. The first column displays the likelihood differences for each 8 combinations of enabling/disabling crossover (COI) and chromatid interference (CI), and zero crossover tetrads (P0). The second column have the estimates for crossover interference parameter *ν* for each chr, whereas the third column depicts the density of the estimated *ν* parameters over all chromosomes with COI/noCI/noP0 (Lep-Rec) and COI/noCI/P0 (gamma and gamma sprinkling). The last column contains the median distance of two crossovers in Lep-Rec, and the density of estimated parameter *p* in the gamma sprinkling models. Chromosomes are in descending length-wise order.

**Figure 10:**
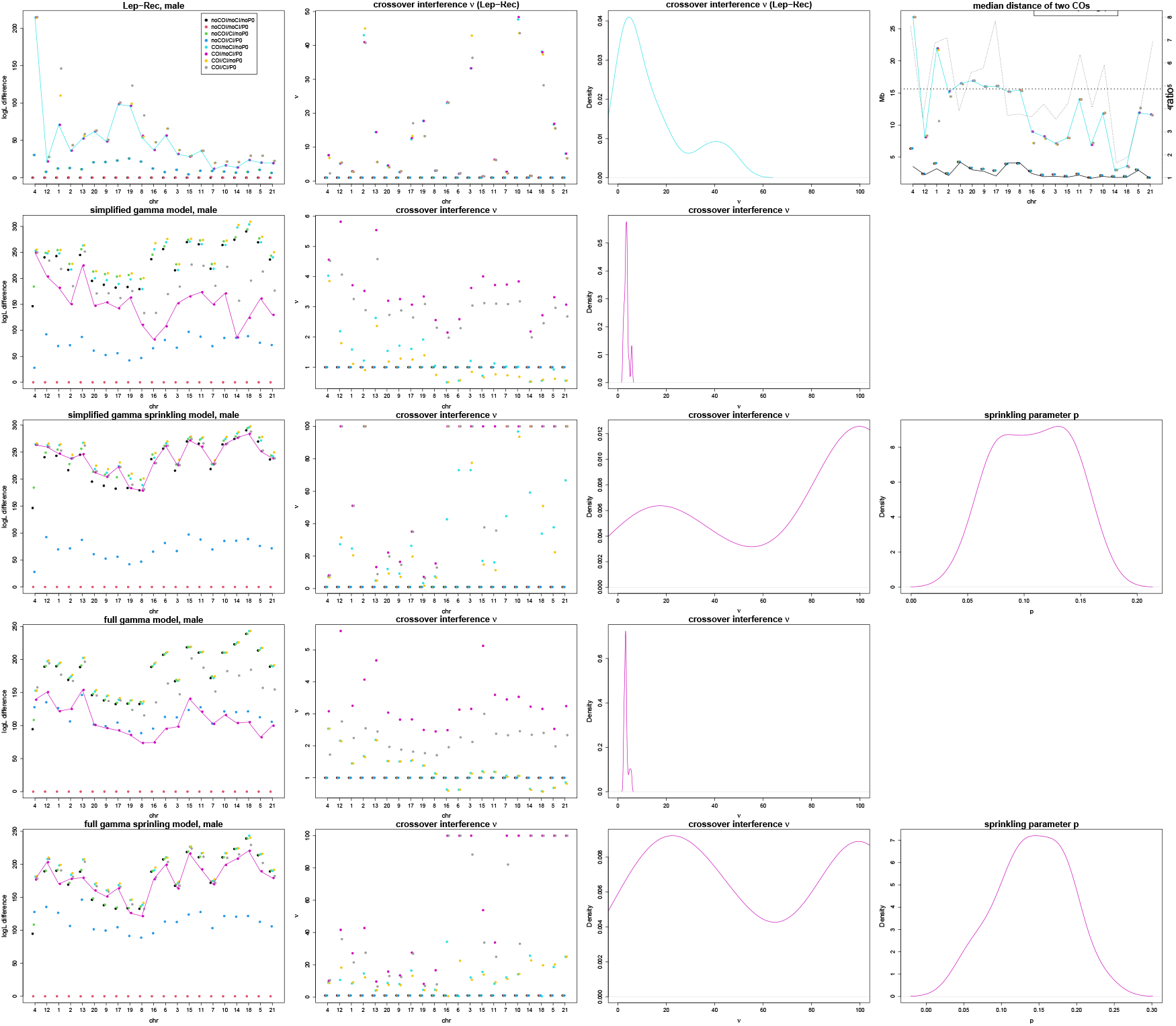
Male fish data for all chromosomes. All rows and columns as in the Figure 9.

**Figure 11:**
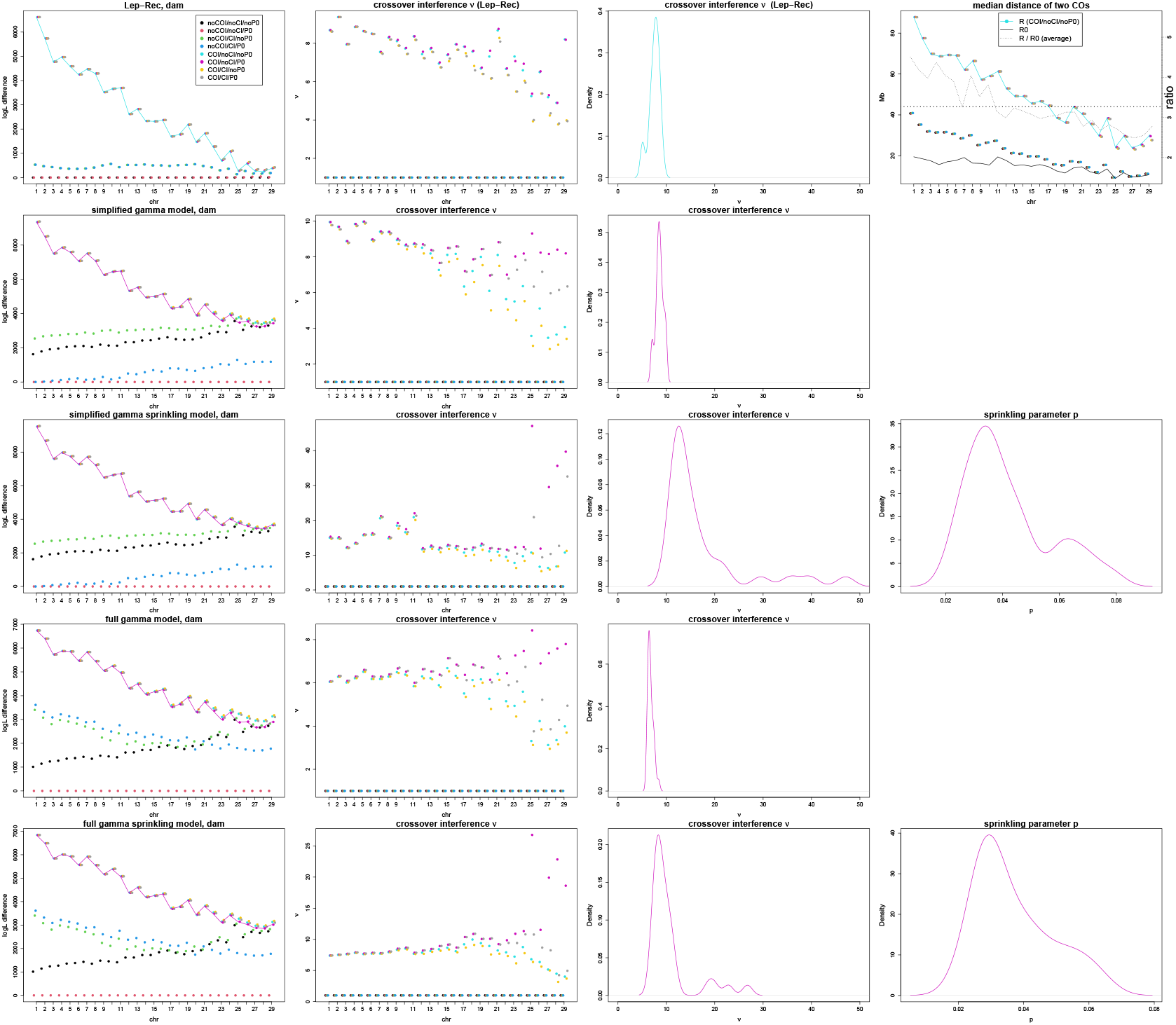
Dam cattle data for all chromosomes. First row is using Lep-Rec, second row is using gamma model and the last gamma sprinkling, all having error modelling enabled. The first column display the likelihood differences for each 8 combinations of enabling/disabling crossover (COI) and chromatid interference (CI), and zero crossover tetrads (P0). The second column have the estimates for crossover interference parameter *ν* for each chr, whereas the third depicts the density of the estimated *ν* parameters over all chromosomes with COI/noCI/noP0 (Lep-Rec) and COI/noCI/P0 (gamma and gamma sprinkling). The last column contains the median distance of two crossovers in Lep-Rec, and the density of estimated parameter *p* in the gamma sprinkling model. Chromosomes are in descending length-wise order.

**Figure 12:**
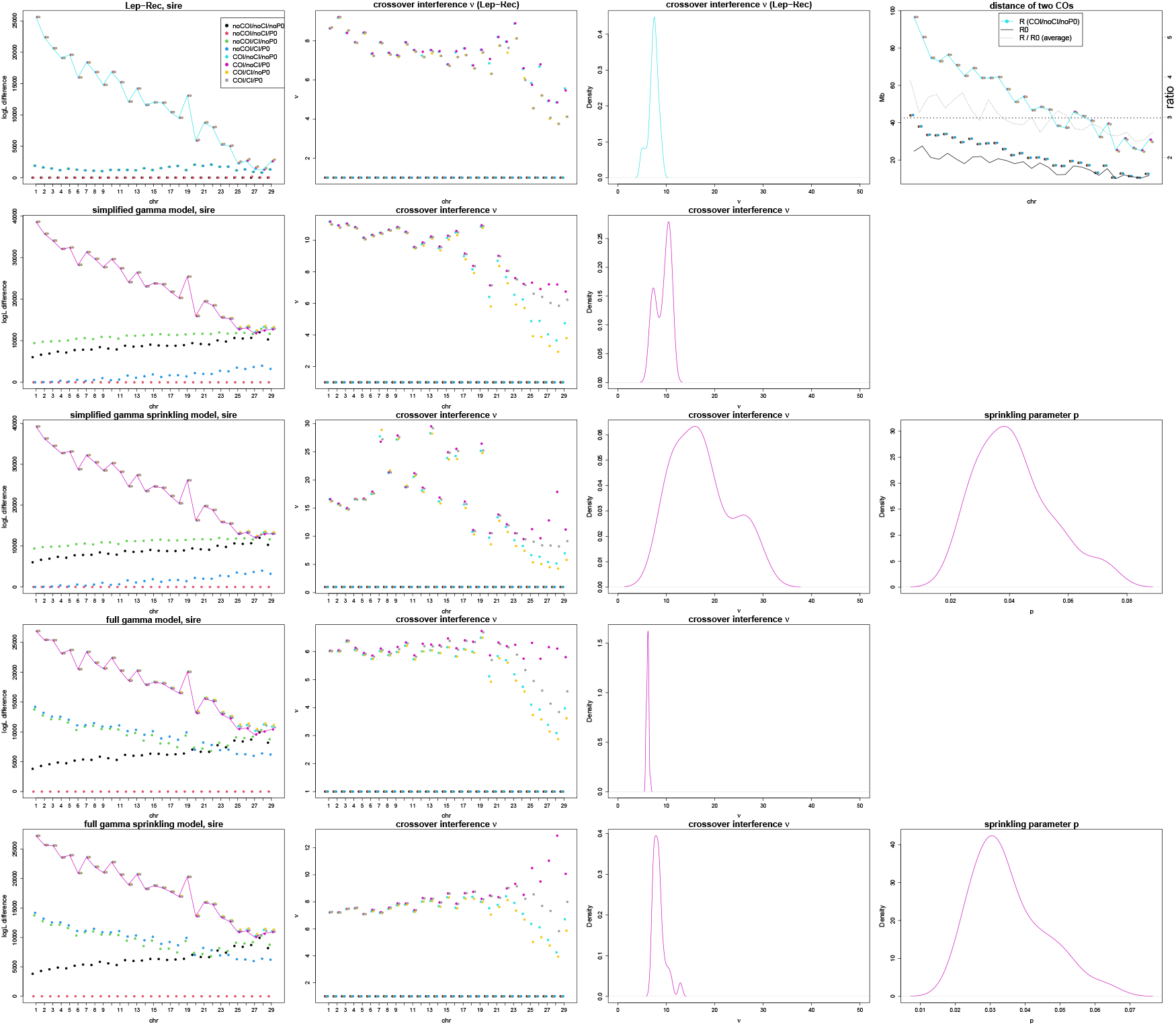
Sire cattle data for all chromosomes. All rows and columns as in the Figure 11.

**Figure 13:**
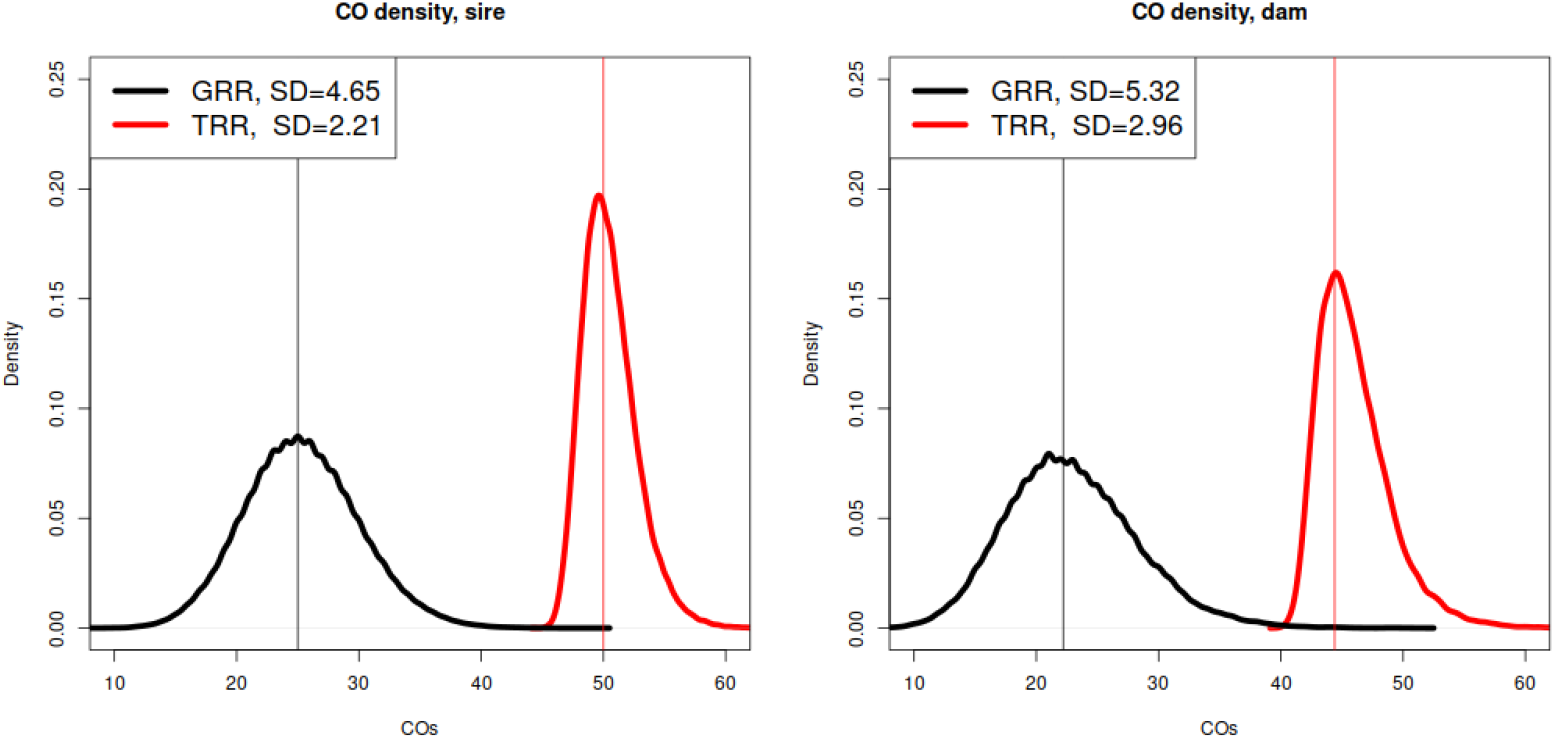
The density of TRR and GRR for the cattle data. The vertical lines are at 50/2 and 50 for sire, and 44.5/2 and 44.5 for dam.

**Figure 14:**
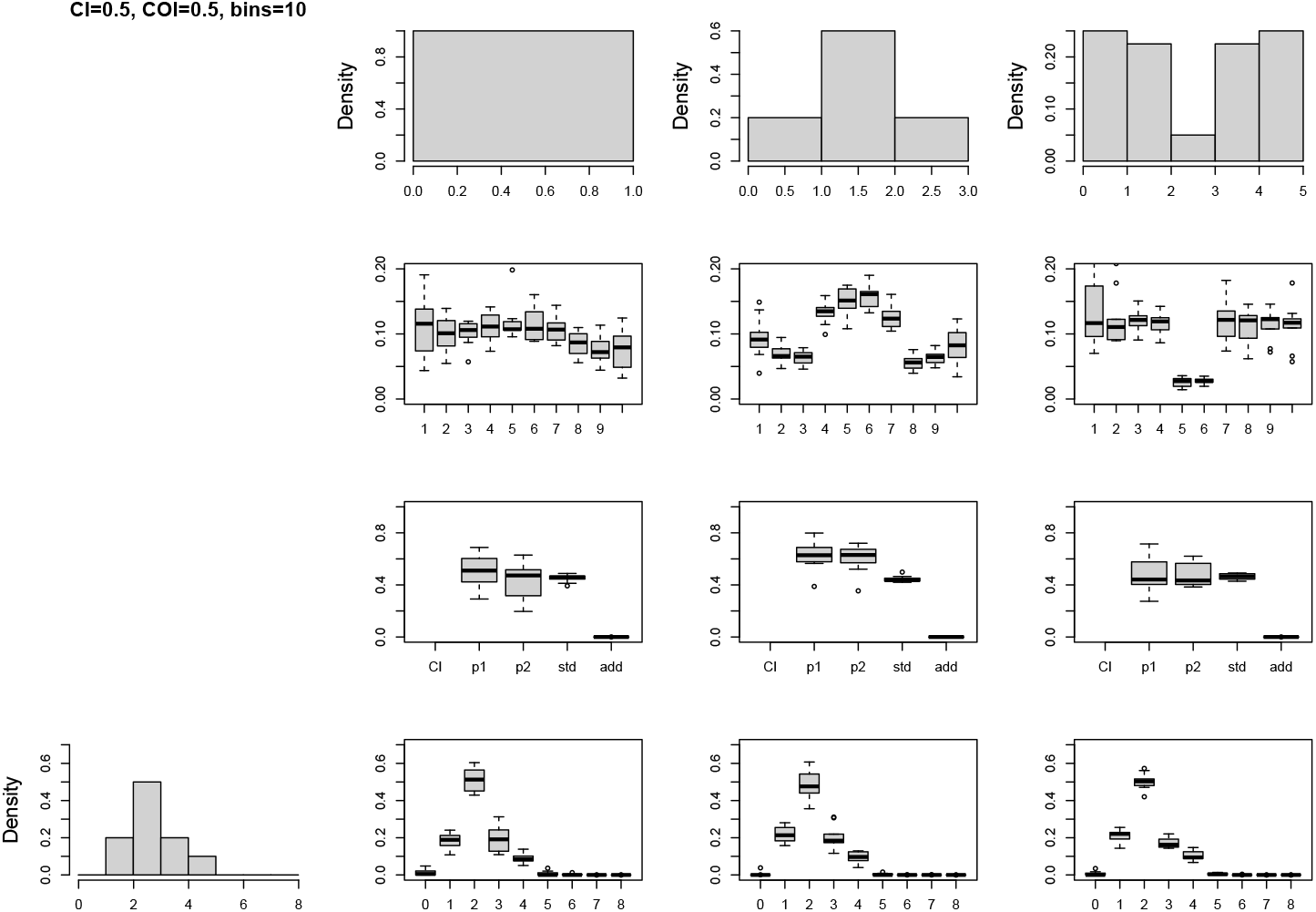
Lep-Rec’s results on simulated data with crossover interference. The bar plots depict the parameter estimates by Lep-Rec over 10 data sets. The histograms on the top display the variable recombination rate used in the simulations, while the leftmost bar plot shows the distribution of tetrad crossovers.

**Figure 15:**
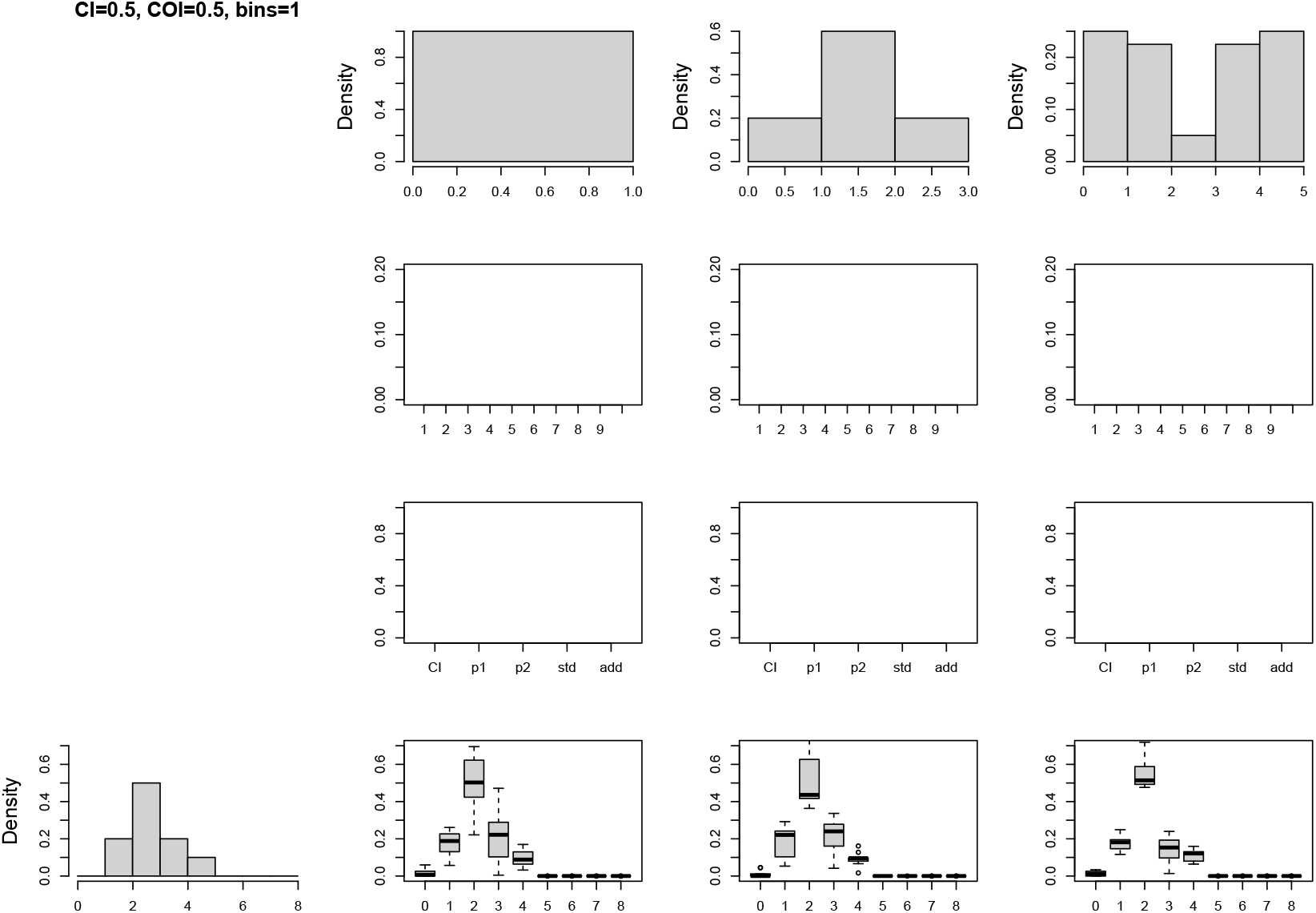
Results on the same data as in Figure 14 with the model of Yu and Feingold (2001) (Lep-Rec with disabled variable recombination and crossover interference, parameter “numBins=1 1”).

**Figure 16:**
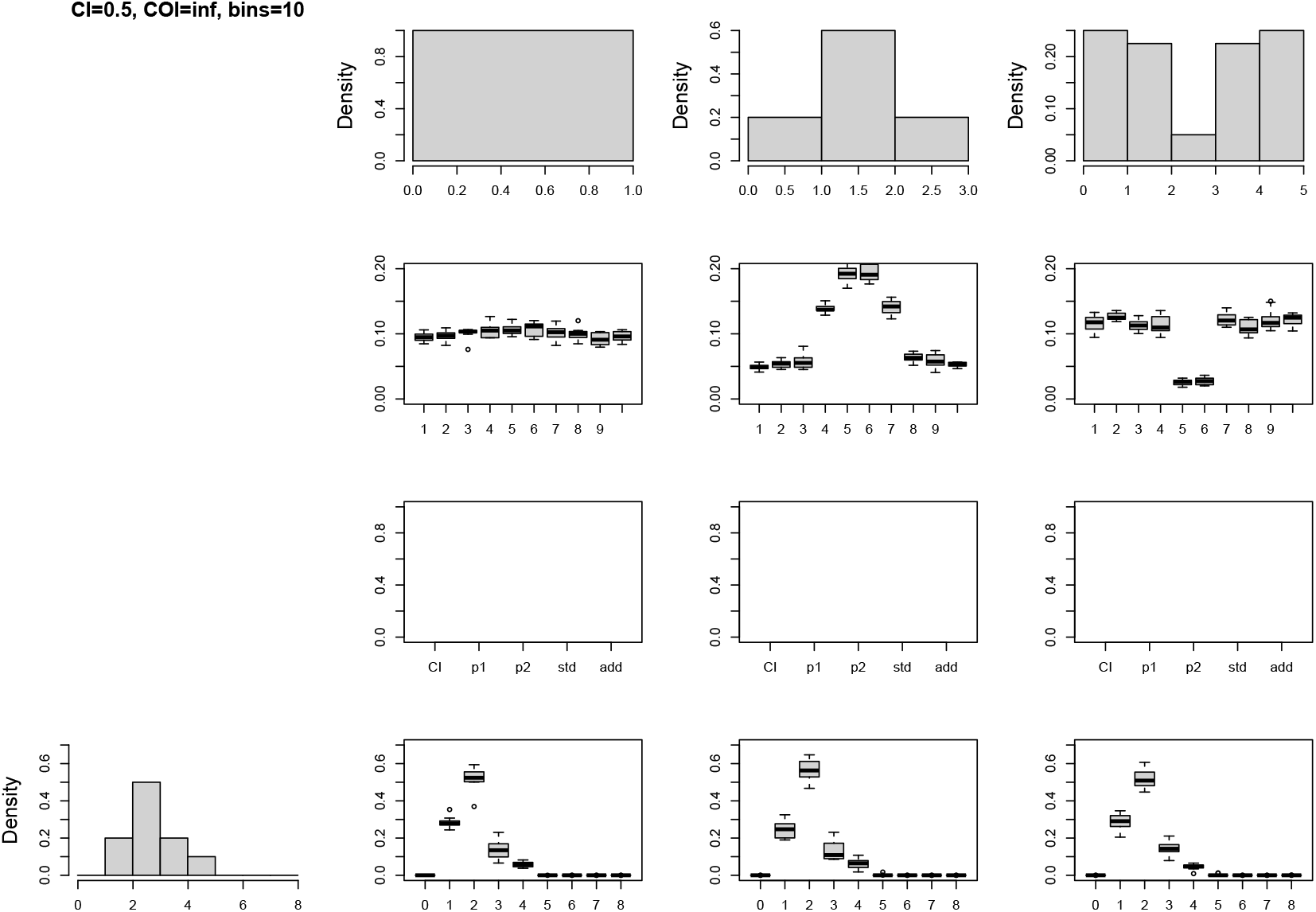
Lep-Rec’s results on simulated data without crossover interference.

**Figure 17:**
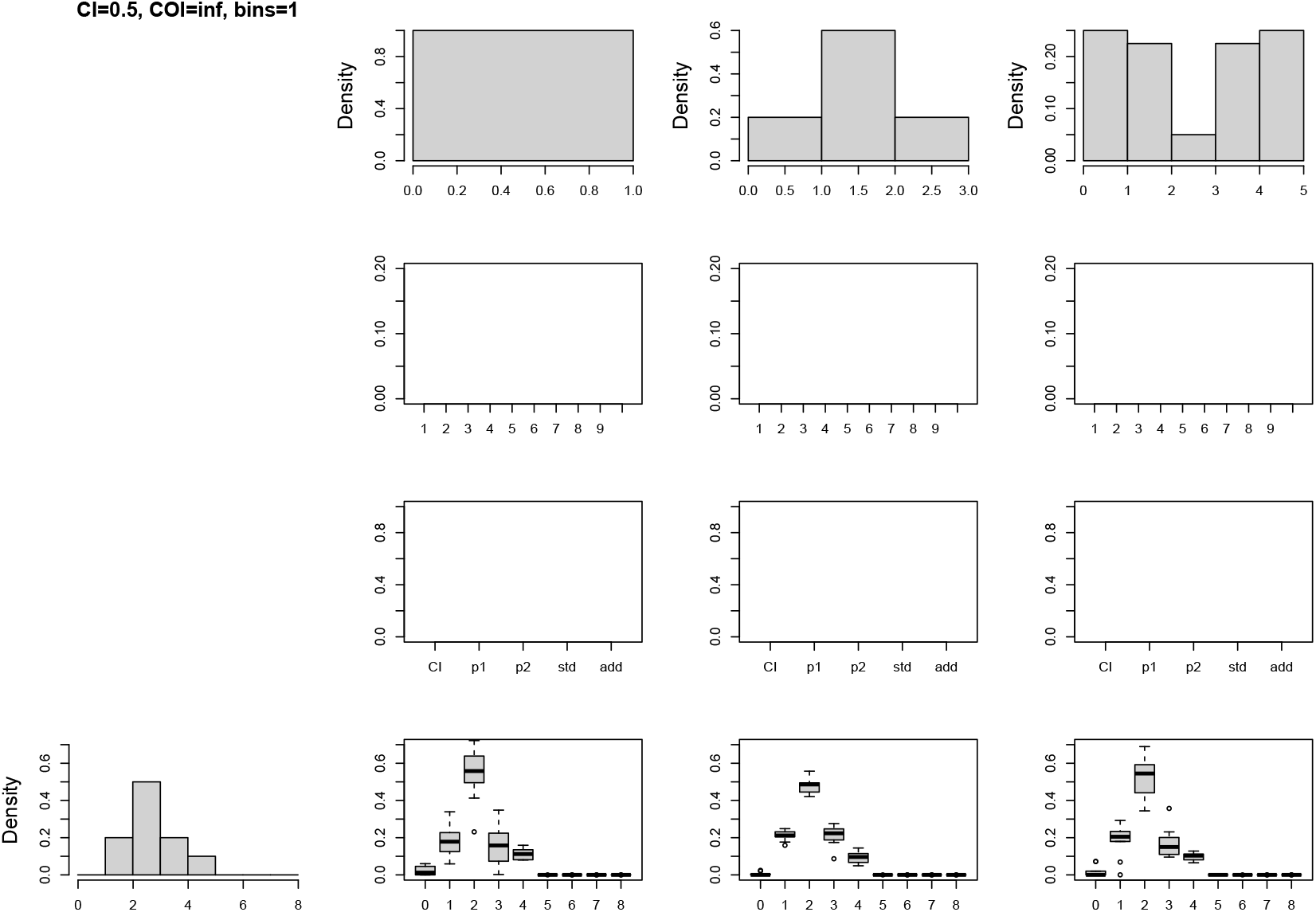
Results on the same data as in Figure 16 with the model of Yu and Feingold (2001).

**Figure 18:**
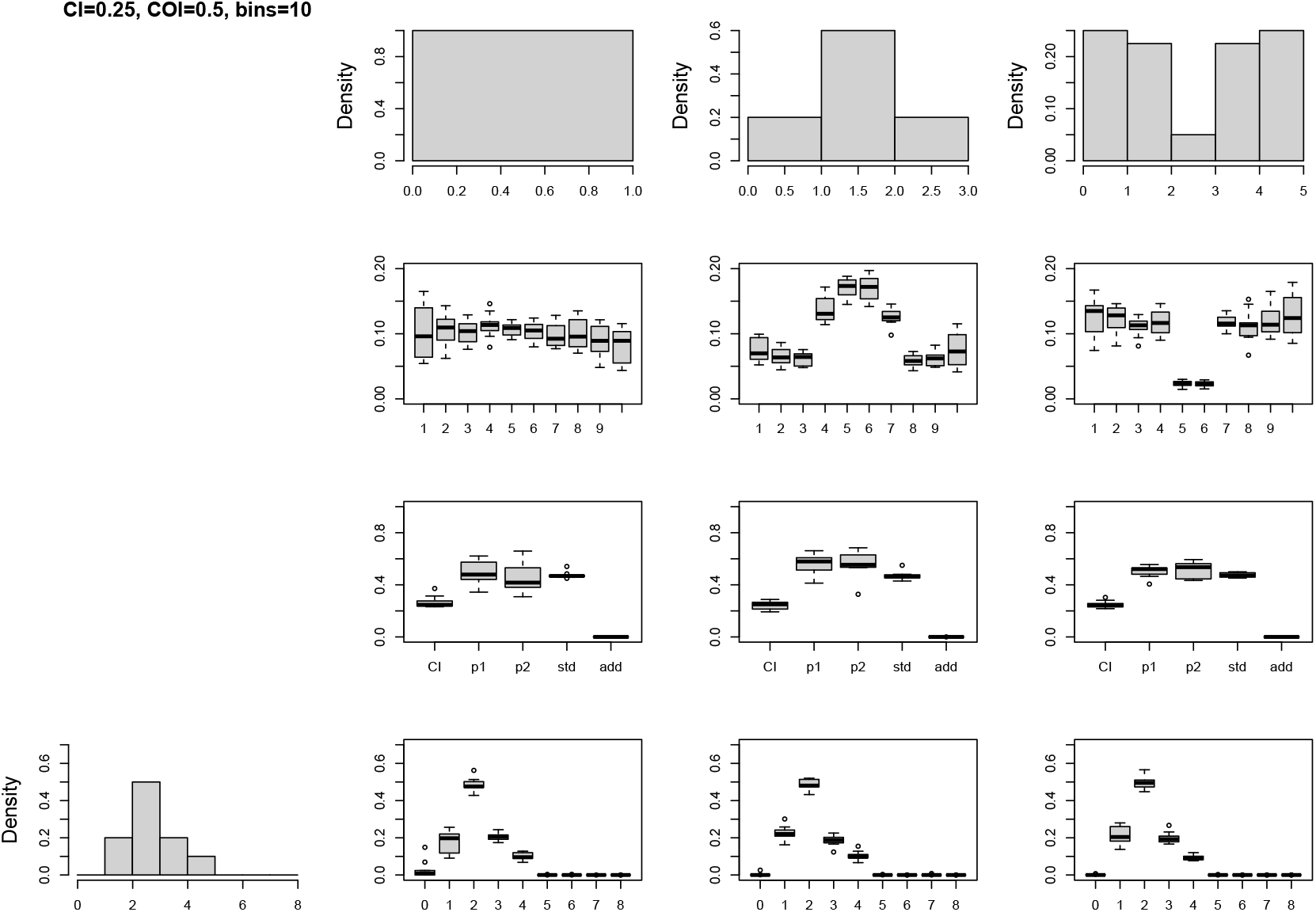
Lep-Rec’s results on simulated data with negative chromatid (*C* = 0.25) and (positive) crossover interference.

**Figure 19:**
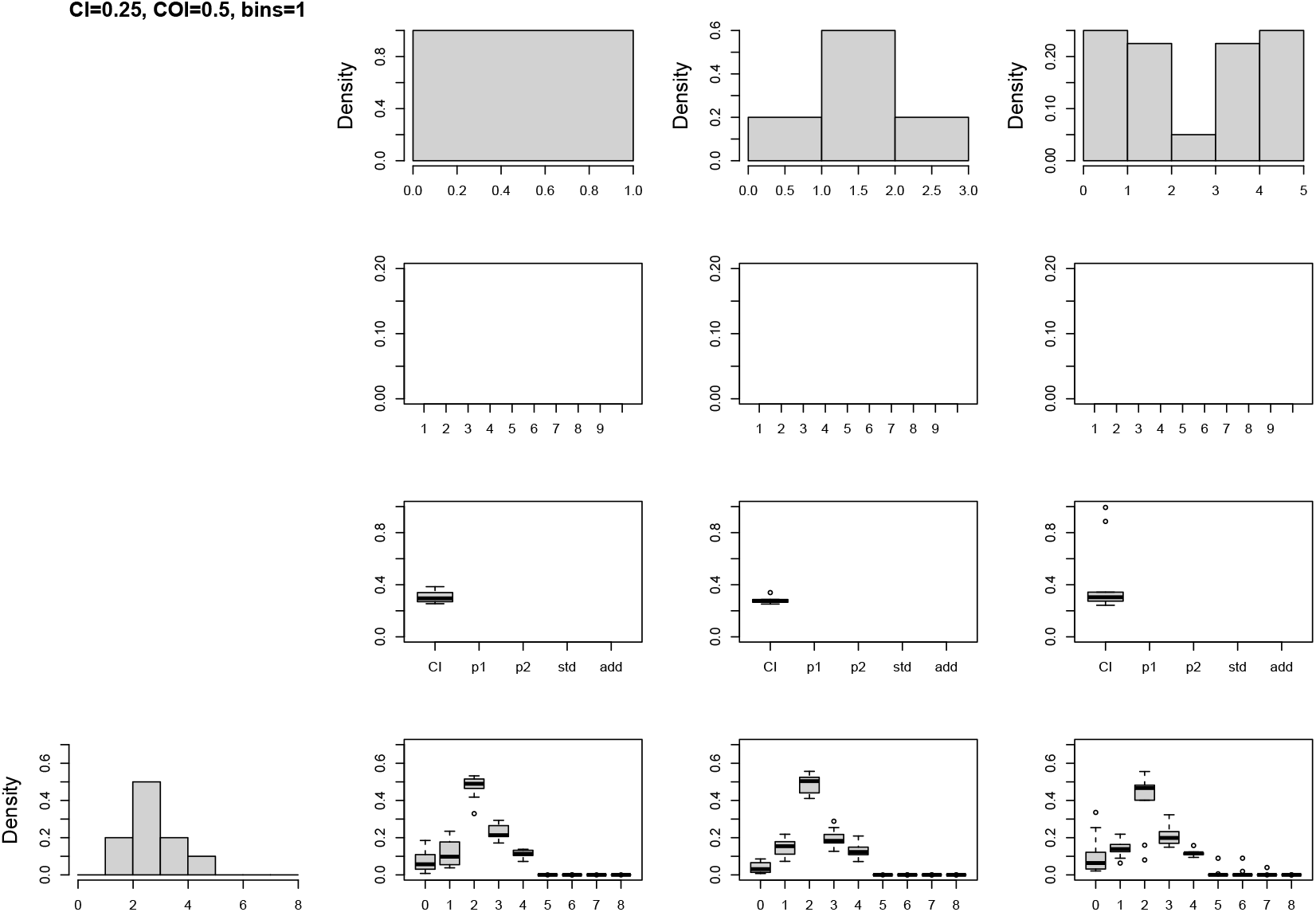
Results on the same data as in Figure 18 with the model of Yu and Feingold (2001) generalised to handle chromatid interference.

**Figure 20:**
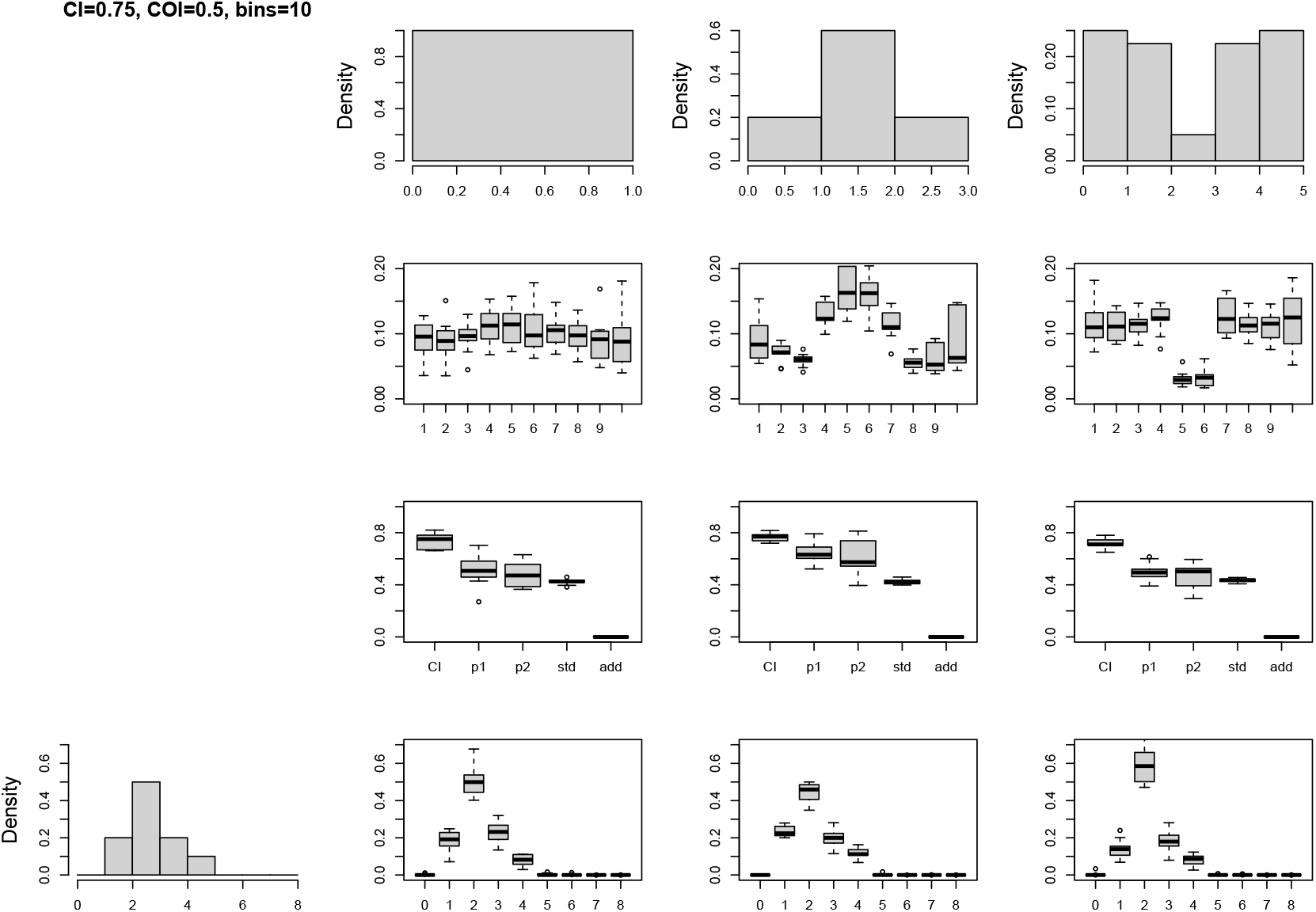
Lep-Rec’s results on simulated data with positive chromatid (*C* = 0.75) and crossover interference.

**Figure 21:**
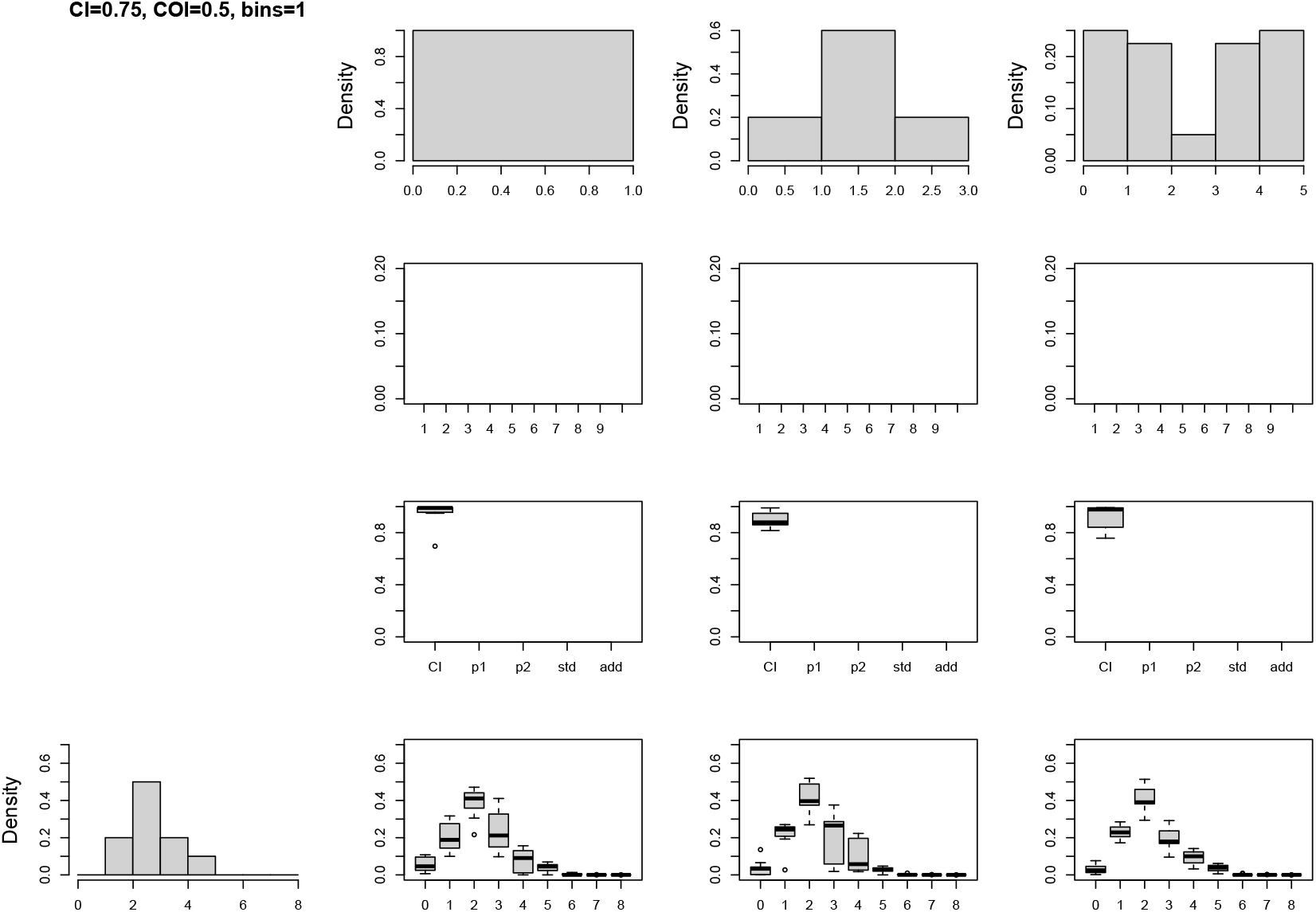
Results on the same data as in Figure 20 with the model of Yu and Feingold (2001) generalised to handle chromatid interference.

## References

Bainbridge, H. E., Brien, M. N., Morochz, C., Salazar, P. A., Rastas, P., and Nadeau, N. J. (2020). Limited genetic parallels underlie convergent evolution of quantitative pattern variation in mimetic butterflies. Journal of evolutionary biology, 33(11):1516–1529.

Brien, M. N., Enciso-Romero, J., Lloyd, V. J., Curran, E. V., Parnell, A. J., Morochz, C., Salazar, P. A., Rastas, P., Zinn, T., and Nadeau, N. J. (2022). The genetic basis of structural colour variation in mimetic Heliconius butterflies. Philosophical Transactions of the Royal Society B, 377(1855):20200505.

Byers, K. J., Darragh, K., Fernanda Garza, S., Abondano Almeida, D., Warren, I. A., Rastas, P. M., Merrill, R. M., Schulz, S., McMillan, W. O., and Jiggins, C. D. (2021). Clustering of loci controlling species differences in male chemical bouquets of sympatric Heliconius butterflies. Ecology and evolution, 11(1):89– 107.

Byers, K. J., Darragh, K., Musgrove, J., Almeida, D. A., Garza, S. F., Warren, I. A., Rastas, P. M., Kučka, M., Chan, Y. F., Merrill, R. M., et al. (2020). A major locus controls a biologically active pheromone component in Heliconius melpomene. Evolution, 74(2):349–364.

Campbell, C. L., Furlotte, N. A., Eriksson, N., Hinds, D., and Auton, A. (2015). Escape from crossover interference increases with maternal age. Nature Communications, 6(1):6260.

Chakravarti, A. (1991). A graphical representation of genetic and physical maps: The marey map. Genomics, 11(1):219–222.

Chen, Z., Xie, L., Tang, X., and Zhang, Z. (2022). Nanocross: A pipeline that detecting recombinant crossover using ont sequencing data. Genomics, 114(6):110499.

Cheng, H., Concepcion, G. T., Feng, X., Zhang, H., and Li, H. (2021). Haplotype-resolved de novo assembly using phased assembly graphs with hifiasm. Nature methods, 18(2):170–175.

Copenhaver, G. P., Housworth, E. A., and Stahl, F. W. (2002). Crossover Interference in Arabidopsis. Genetics, 160(4):1631–1639.

Darragh, K., Orteu, A., Black, D., Byers, K. J., Szczerbowski, D., Warren, I. A., Rastas, P., Pinharanda, A., Davey, J. W., Fernanda Garza, S., et al. (2021). A novel terpene synthase controls differences in anti-aphrodisiac pheromone production between closely related Heliconius butterflies. PLoS Biology, 19(1):e3001022.

Davey, J. W., Barker, S. L., Rastas, P. M., Pinharanda, A., Martin, S. H., Durbin, R., McMillan, W. O., Merrill, R. M., and Jiggins, C. D. (2017). No evidence for maintenance of a sympatric Heliconius species barrier by chromosomal inversions. Evolution letters, 1(3):138–154.

Davey, J. W., Chouteau, M., Barker, S. L., Maroja, L., Baxter, S. W., Simpson, F., Merrill, R. M., Joron, M., Mallet, J., Dasmahapatra, K. K., et al. (2016). Major improvements to the Heliconius melpomene genome assembly used to confirm 10 chromosome fusion events in 6 million years of butterfly evolution. G3: Genes, Genomes, Genetics, 6(3):695–708.

De Lorenzi, L., Iannuzzi, A., Rossi, E., Bonacina, S., and Parma, P. (2017). Centromere repositioning in cattle (bos taurus) chromosome 17. Cytogenet. Genome Res., 151(4):191–197.

Fisher, R. (1925). Statistical methods for research workers. Edinburgh Oliver & Boyd.

Gauthier, F., Martin, O. C., and Falque, M. (2011). Coda (crossover distribution analyzer): quantitative characterization of crossover position patterns along chromosomes. BMC Bioinformatics, 12:27–27.

Housworth, E. and Stahl, F. (2003). Crossover interference in humans. The American Journal of Human Genetics, 73(1):188–197.

Kadri, N. K., Harland, C., Faux, P., Cambisano, N., Karim, L., Coppieters, W., Fritz, S., Mullaart, E., Baurain, D., Boichard, D., Spelman, R., Charlier, C., Georges, M., and Druet, T. (2016). Coding and noncoding variants in hfm1, mlh3, msh4, msh5, rnf212, and rnf212b affect recombination rate in cattle. Genome Research, 26(10):1323–1332.

Karlin, S. and Liberman, U. (1979). A natural class of multilocus recombination processes and related measures of crossover interference. Advances in Applied Probability, 11(3):479–501.

Kemppainen, P., Li, Z., Rastas, P., Löytynoja, A., Fang, B., Yang, J., Guo, B., Shikano, T., and Merilä, J. (2021). Genetic population structure constrains local adaptation in sticklebacks. Molecular Ecology, 30(9):1946–1961.

Kivikoski, M., Rastas, P., Löytynoja, A., and Merilä, J. (2021a). Automated improvement of stickleback reference genome assemblies with lep-anchor software. Molecular Ecology Resources, 21(6):2166–2176.

Kivikoski, M., Rastas, P., Löytynoja, A., and Merilä, J. (2021b). Automated improvement of stickleback reference genome assemblies with lep-anchor software. Molecular Ecology Resources, 21(6):2166–2176.

Li, H. (2013). Aligning sequence reads, clone sequences and assembly contigs with bwa-mem.

McPeek, M. and Speed, T. (1995). Modeling interference in genetic recombination. Genetics, 139(2):1031–44.

Ott, J. (1996). Estimating crossover frequencies and testing for numerical interference with highly polymorphic markers. In Speed, T. and Waterman, M. S., editors, Genetic Mapping and DNA Sequencing, pages 49–63, New York, NY. Springer New York.

Otto, S. P. and Payseur, B. A. (2019). Crossover interference: Shedding light on the evolution of recombination. Annual Review of Genetics, 53(1):19–44.

Peñalba, J. V. and Wolf, J. B. W. (2020). From molecules to populations: appreciating and estimating recombination rate variation. Nature Reviews Genetics, 21(8):476–492.

Rastas, P. (2017). Lep-MAP3: robust linkage mapping even for low-coverage whole genome sequencing data. Bioinformatics, 33(23):3726–3732.

Rastas, P. (2020). Lep-Anchor: automated construction of linkage map anchored haploid genomes. Bioinformatics, 36(8):2359–2364.

Rezvoy, C., Charif, D., Guéguen, L., and Marais, G. A. (2007). MareyMap: an R-based tool with graphical interface for estimating recombination rates. Bioinformatics, 23(16):2188–2189.

Sardell, J. M. and Kirkpatrick, M. (2020). Sex differences in the recombination landscape. The American Naturalist, 195(2):361–379.

Sarens, M., Copenhaver, G., and De Storme, N. (2021). The role of chromatid interference in determining meiotic crossover patterns. Front. Plant Sci., 12:656691.

Siberchicot, A., Bessy, A., Guéguen, L., and Marais, G. A. (2017). MareyMap Online: A User-Friendly Web Application and Database Service for Estimating Recombination Rates Using Physical and Genetic Maps. Genome Biology and Evolution, 9(10):2506–2509.

Stapley, J., Feulner, P. G., Johnston, S. E., Santure, A. W., and Smadja, C. M. (2017). Variation in recombination frequency and distribution across eukaryotes: patterns and processes. Philosophical Transactions of the Royal Society B: Biological Sciences, 372(1736):20160455.

Sturtevant, A. H. (1913). The linear arrangement of six sex-linked factors in drosophila, as shown by their mode of association. Journal of Experimental Zoology, 14(1):43–59.

Suomalainen, E. (1966). Achiasmatische oogenese bei trichopteren. Chromosoma, 18(2):201–207.

Terrell, G. R. and Scott, D. W. (1992). Variable kernel density estimation. The Annals of Statistics, 20(3):1236–1265.

The Heliconius Genome Consortium (2012). Butterfly genome reveals promiscuous exchange of mimicry adaptations among species. Nature, 487(7405):94–98.

Tibshirani, R. J. and Efron, B. (1993). An introduction to the bootstrap. Monographs on statistics and applied probability, 57(1):1–436.

Wang, S., Zickler, D., Kleckner, N., and Zhang, L. (2015). Meiotic crossover patterns: Obligatory crossover, interference and homeostasis in a single process. Cell Cycle, 14(3):305–314.

Weinstein, A. (1936). The theory of multiple-strand crossing over. Genetics, 21(3):155–199.

Yu, K. and Feingold, E. (2001). Estimating the frequency distribution of crossovers during meiosis from recombination data. Biometrics, 57(2):427– 434.

Zhao, H., McPeek, M. S., and Speed, T. P. (1995). Statistical analysis of chromatid interference. Genetics, 139(2):1057–1065.

